# A neuronal CRISPRi screen identifies PQLC2 as a lysosomal pH regulator controlling tau homeostasis

**DOI:** 10.64898/2026.08.25.747102

**Authors:** Mackenzie Welch, Paul J. Sampognaro, Shunpan Shu, Kriti Chaplot, Anuja Bothra, Patricia A. Castruita, Andrew W. Smith, Tara Antee, Molly Hodul, Ruilin Tian, Vienna Gao, Juanita C. Limas, Kevin D. Burris, Joanne L. Parker, Jennifer S. Yokoyama, Bruce L. Miller, William W. Seeley, Simon Newstead, Martin Kampmann, Aimee W. Kao

## Abstract

Lysosomes make key contributions to the maintenance of cellular proteostasis, and their functional compromise has been linked to aging and neurodegenerative disease. A defining characteristic of lysosomes is their relative acidity compared to other subcellular compartments, a quality that enables the efficient breakdown of macromolecules. Evidence suggests that neuronal lysosomal pH becomes dysregulated with aging and neurodegenerative disease, yet the mechanisms by which lysosomal pH is maintained remain incompletely understood. To better understand neuronal lysosomal pH regulation, we conducted a genome-wide CRISPRi-based screen in iPSC-derived iNeurons for modifiers of lysosomal pH. We validated several previously known regulators of lysosomal pH and identified novel pathways capable of modifying lysosomal pH, including protein UFMylation and mitochondrial homeostasis. We demonstrate that loss of the lysosomal cationic amino acid exporter, PQLC2, prevents lysosomal acidification in a manner independent of amino acid transport. A novel, tauopathy-associated mutation in *PQLC2* impairs lysosomal acidification and drives tau accumulation. Together, this study reveals novel genes that modify lysosomal pH and highlights potential new targets for ameliorating age-related lysosome dysfunction.

## Introduction

Lysosomes are acidic, membrane-bound organelles that serve as a major site of protein degradation within the cell^1^. Beyond protein degradation, lysosomes play central roles in autophagy regulation^2^, nutrient sensing^3^, and calcium signaling^4^. Optimal lysosomal function relies heavily on maintenance of an acidic environment within the lysosome, with resident acid hydrolases tuned to function optimally at pH ∼4.5-5.0^5^. This pH dependence reflects the sensitivity of protein structure to even relatively small changes in pH^6^. Lysosomal acidification requires coordination of multiple processes: the assembly of the vacuolar-type H^+^-ATPase (V-ATPase) on the lysosomal membrane^7^, the import of protons by the V-ATPase^8^, and the offset of positively charged protons by counter-ion currents from other lysosomal membrane channels, including MCOLN1, TMEM175, and CLC-7^9–11^. Without the proper action of these channels and transporters, lysosomal pH can drift and impact the function of resident lysosomal hydrolases. Even modest changes in lysosomal pH can also alter substrate specificity, including that of specific proteases for neurodegenerative proteins such as alpha-synuclein, TDP-43, and tau^12,13^. Thus, proper lysosomal pH maintenance is likely key to optimal proteostasis and the prevention of protein aggregation disorders.

Given the importance of lysosomal pH maintenance to proteostasis, it is not surprising that age-associated loss of lysosomal acidity has been observed in both *S. cerevisiae* and *C. elegans*^14,15^. Genetic models of neurodegenerative diseases have also demonstrated impairments in lysosomal pH homeostasis^16–19^. Notably, recent work has shown lysosomal pH alkalinization precedes cell death and amyloid beta deposition in a mouse model of Alzheimer’s disease^20^. Restoring lysosomal pH homeostasis reduces the accumulation of alpha-synuclein in cell models of Parkinson’s disease^10,21,22^. As aging represents the primary shared risk factor for all neurodegenerative disease^23^, lysosomal pH dysregulation could serve as a key potential link between aging and neurodegeneration.

Systematic approaches to understanding how neuronal lysosomal pH is maintained are lacking. This is partly due to a lack of appropriate tools for large-scale experiments involving lysosomal pH. Previous methods for measuring lysosomal pH have relied on pH-sensitive fluorescent molecules such as Lysosensor or Oregon Green 488, either as free dyes or dextran-conjugates^24^. These methods present a challenge for large studies that include genetic screens, primarily due to challenges in delivery, cost, and potential for lysosomal pH disruption by the dye itself^25^. To overcome these limitations, we recently developed a genetically encoded lysosomal pH sensor, Fluorescence Indicator REporting pH in Lysosomes (FIRE-pHLy)^26^. As a genetically encoded, ratiometric reporter of lysosomal pH, FIRE-pHLy is amenable to high-throughput applications as we have utilized it for screening to identify novel compounds capable of acidifying lysosomes^27^.

To better understand the genetic and pathway regulators, we performed a genome-wide CRISPR interference (CRISPRi) screen for modifiers of lysosomal pH in human induced pluripotent stem cell (iPSC)-derived neurons (iNeurons). We identified multiple disease genes as well as numerous pathways not previously linked to lysosomal pH, including protein UFMylation and the mitochondrial ribosome. Notably, the cationic amino acid exporter PQLC2 emerged as one of the strongest lysosome-localized hits. Although PQLC2 is increasingly recognized as a key regulator of lysosomal pH homeostasis^28^, its role in maintaining physiological lysosomal pH in neurons has remained unexplored. Additionally, we find that PQLC2 serves as a regulator of tau homeostasis. These results elucidate novel genes and pathways underpinning lysosomal pH homeostasis, advance understanding of direct and indirect effects on lysosomal pH setpoints and identify a novel genetic risk factor for tauopathy.

## Results

### Genome-wide screen for modifiers of lysosomal pH in iNeurons

Despite the critical importance of lysosomal pH in aging and neurodegeneration, we lack a comprehensive understanding of the genes regulating lysosomal pH homeostasis in neurons. To address this gap, we utilized the genetically encoded lysosomal pH sensor, FIRE-pHLy, in a CRISPRi screen for modifiers of lysosomal pH in human iNeurons. FIRE-pHLy consists of a luminal-facing pH-sensitive fluorophore, mTFP, fused to lysosomal-associated membrane protein-1 (LAMP1) for lysosome targeting and a cytosol-facing mCherry for ratiometric measurement (**Fig. 1a)**^26^. We stably transduced doxycycline-inducible neurogenin-2 (NGN2) iPSCs containing CRISPRi machinery^29^ with lentivirus expressing a FIRE-pHLy construct. The ratio of mTFP:mCherry increased with bafilomycin, validating the sensor function in iNeurons (**Fig. 1b)**. To ensure that FIRE-pHLy was properly trafficked to lysosomes in iNeurons, we measured colocalization between the sensor and the lysosome marker LAMP2. FIRE-pHLy exhibited significantly higher colocalization with LAMP2 compared to mitochondria and early endosomes (**Fig. 1c,d**). FIRE-pHLy iPSCs were then tested to ensure that iNeurons were amenable to Fluorescence-Activated Cell Sorting (FACs) sorting and responsive to changes in lysosomal pH (**Extended Data Fig. 1a,b)**.

**Fig. 1:**
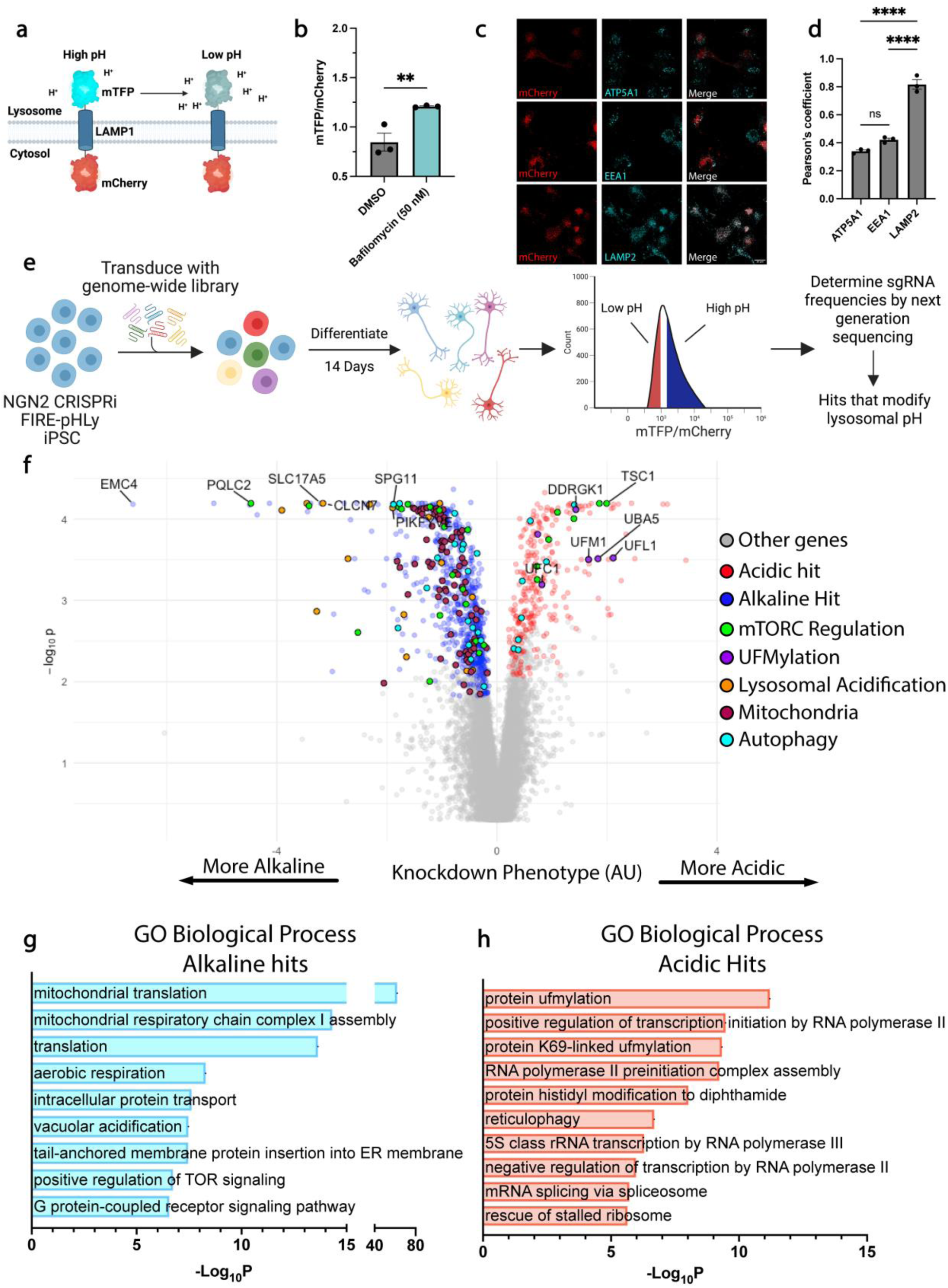
Whole genome screen for modifiers of lysosomal pH in human iNeurons. **a**, Schematic of genetically encoded lysosomal pH reporter, FIRE-pHLy. **b**, Overnight incubation with 50 nM bafilomycin-A1 increases mTFP:mCherry ratio, demonstrating alkalinization of lysosomal pH. **c**, Representative images of neuronally expressed FIREpHLy colocalized with LAMP2-positive organelles compared to EEA1 and ATP5A1. **d**, Pearson’s r correlations of mCherry with LAMP2, EEA1 or ATP5A1 (n=1-2 wells imaged from 3 independent replicates) demonstrating strong colocalization with lysosomes. **e**, Schematic of iNeuron lysosomal pH screen. **f,** Volcano plot of hit genes from genome-wide screen. Phenotype (normalized log2 ratio of counts in the mTFP:mCherry high versus mTFP:mCherry low populations) is plotted versus the negative log10 of the p-value, calculated with a Mann-Whitney U-test. Acidic hits are in red, and alkaline hits are in blue. **g**, GO biological process terms for Alkaline pH hits. **h**, GO biological process terms for acidic pH hits. * p <0.05, ** p <0.01, *** p <0.001.

We then performed a whole-genome screen for modifiers of lysosomal pH in iNeurons (**Fig. 1e**). After sample processing, we identified 1,051 genes that modified lysosomal pH in iNeurons (**Fig. 1f, Table S1**). Knockdown of three-quarters of the hit genes had an alkaline lysosomal pH phenotype with the remaining hits resulting in an acidic phenotype. For clarity, genes whose CRISPRi knock-down result in an alkaline phenotype will be referred to as alkalinizing hits and those that have an acidic phenotype when knocked-down will be referred to as acidifying hits.

Among the acidifying hits, several Gene Ontology (GO) categories of interest were identified. Every member of the UFM1 conjugation pathway (*UFM1*, *UFL1*, *UBA5*, *DDRGK1*, *CDK5RAP3* and *UFC1*) was enriched within the acidifying portion of the screen (**Fig. 1f,h**). Protein UFMylation is a ubiquitin-like process recently identified to be a modifier of tau homeostasis^30,31^. Knockdown of members of the UFMylation pathway reduced the amount of both tau aggregation and tau oligomers, and the corresponding decrease in lysosomal pH in our model links lysosomal pH changes with tau homeostasis. Previous work has linked deficiency of DDRGK1 to autophagy induction with dysfunctional lysosomes^32^, indicating that the acidic phenotype may be due to compensatory hyper-acidification of lysosomes. Multiple additional GO categories refer to transcriptional and translational processes, also supporting compensatory acidification of lysosomes in these hits (**Fig. 1h**).

Because their knock-down led to an alkaline phenotype, alkalinizing hits are more likely to represent genes whose normal function is to contribute to neuronal lysosomal acidification. In support of this, the GO enrichment analysis of alkalinizing hits identified vacuolar acidification and positive regulation of mTOR signaling (**Fig. 1g)**^27,33^. Supporting the biological relevance of this screen, knockdown of known lysosomal pH acidifiers *CLCN7*, *OSTM1, TMEM106B, MCOLN1, PIKFYVE,* and multiple v-ATPase subunits all resulted in an alkaline pH phenotype^11,34–36^. Interestingly, several neurodegenerative disease genes were found to modify lysosomal pH in this dataset, including *TMEM106B*, *ATP13A2*, *MCOLN1*, *DNAJC13* and *SCARB2,* underscoring the association between neurodegeneration and lysosomal homeostasis.

The v-ATPase consumes energy to pump protons into the lysosome; thus, we speculated that genes whose knockdown would disrupt energy metabolism would be found among the alkalinizing hits. Consistent with this, along with V-ATPase subunits, alkalinizing hits identified mitochondrial translation, mitochondrial complex I assembly and respiration as enriched categories (**Fig. 1g**). When compared to a previous CRISPRi screen for modifiers of ATP levels in iNeurons^37^, alkalinizing hits largely corresponded to genes that decreased cellular ATP levels (**Extended Data Fig. 2a**). Importantly, many of the highest-performing hits from our screen formed a connected network, with nodes centered around mitochondrial ATP generation, V-ATPase function, mTORC regulation, and UFMylation (**Extended Data Fig. 2b**).

### Secondary screens validate FIRE-pHLy as a robust lysosomal pH reporter

To validate the whole-genome screen results, we constructed a lysosomal pH sgRNA sub-library comprising 212 control sgRNAs and 4665 sgRNAs targeting 831 of the highest-scoring hits from the whole-genome screen, genes annotated as lysosomal resident proteins and/or linked to neurodegenerative disease (**Table S3)**. We first repeated our FIRE-pHLy screen with the sub-library to validate our whole genome lysosomal pH screen; this largely correlated the primary screen hits (**Fig. 2b, Table S1**).

**Figure 2:**
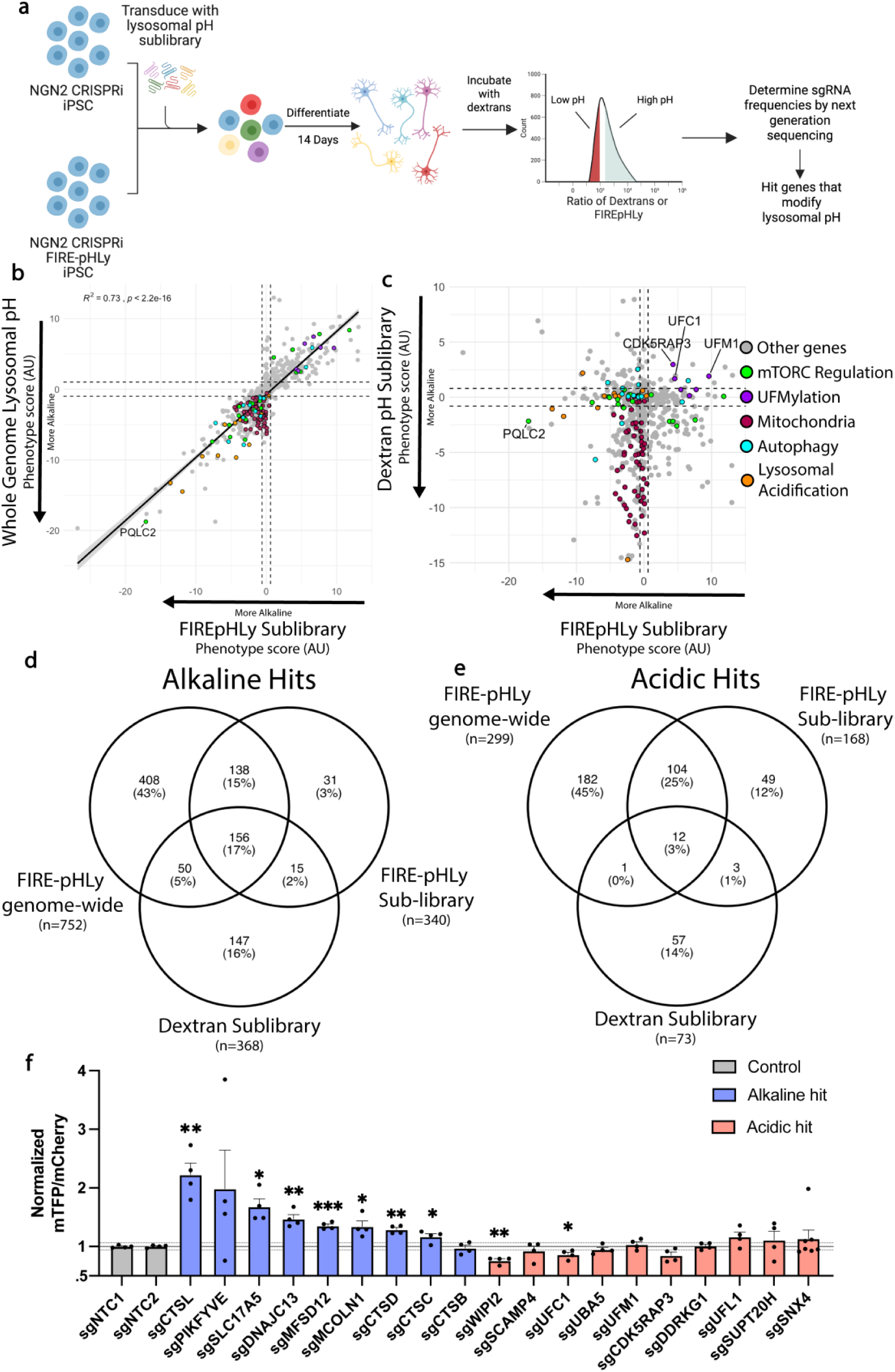
Secondary validation of CRISPRi screen for modifiers of lysosomal pH. **a**, Sublibrary iNeuron screen schematic. Cells expressing FIRE-pHLy or no sensor were transduced with the pH sgRNA sublibrary and differentiated into neurons for two weeks. For dextran-based lysosomal pH measurement, cells were stained with Oregon Green 488-dextran and CF555-Dextran overnight and chased to the lysosome with a 4-hour incubation in fresh media. Cells with either FIRE-pHLy or the dextrans were dissociated and high and low lysosome pH fractions were collected by FACS. **b**, Correlation between sub-library FIRE-pHLy screen phenotype score and whole genome screen phenotype score (Pearson’s r = 0.73). **c,** Comparison between FIRE-pHLy phenotype score and dextran-based sublibrary screen phenotype score. d, Overlap between alkalinizing hit genes in whole genome and two sub-library screens. **e,** Overlap between acidic hit genes in whole genome and two sub-library screens. **f**, High-content imaging validation of select hit genes using individually cloned sgRNAs analyzed by one-way ANOVA. Mean of 4 independent replicates, blue bars were alkaline hits, and red bars were acidic hits in our initial screen. Error bars represent standard error. Dashed line is threefold standard deviation of the non-targeting control mTFP:mCherry ratios. * p <0.05, ** p <0.01, *** p <0.001.

We then used the sub-library to compare FIRE-pHLy against an orthogonal measure of lysosomal pH, a dextran-based method using a combination of pH-sensitive and insensitive dyes (**Fig. 2a,c**)^22^. In comparing the FIRE-pHLy and dextran-based pH secondary screens, we noted good concordance between the mitochondrial and UFMylation pathway hits, and both screens identified more alkalinizing than acidifying hits (**Fig. 2c-e**). Notably, the mTORC signaling and autophagy pathway hits showed limited correlation between the two screens and even included genes with opposing phenotypes. Within the genes that held opposing phenotypes, many are known to regulate either endosomal transport or cellular protein trafficking including *WDR91*, *GNPTAB*, *RAB3GAP1* and *RAB1A*^38–40^. FIB-SEM analysis has shown that dextran-based lysosomal pH sensors can fail to reach 20-25% of lysosomes^41^, which likely explains the differences between the dextran-based probe results and those from our genetically encoded FIRE-pHLy. Importantly, several of the genes previously associated with lysosomal acidification were hits within our FIRE-pHLy screen, and not with the dextran-based method, demonstrating the ability of FIRE-pHLy to report lysosomal pH in high-throughput applications.

Finally, we generated stable cell lines from a subset of hits of interest from our FIRE-pHLy screen and tested these manually by high-content confocal imaging. This approach validated robust increases in lysosomal pH for alkalinizing targets, as nearly all recapitulated (**Fig. 2f**). In contrast, several of the acidic hits did not validate. This may be because acidic hits result from compensatory responses to impairments in proteostasis machinery, which is supported by the GO terms (**Fig. 1h**). It may also suggest a simple challenge of lysosomal pH screening—the narrow dynamic range in which to measure acidification. Lysosomal proton leak channels and other transporters protect against over-acidification^10^. Nonetheless, the acidic hits could represent important targets for therapeutics aimed at lysosomal acidification.

### PQLC2 regulates lysosomal pH and alters lysosomal function in human iNeurons

Given our interest in identifying proteins that directly contribute to neuronal lysosomal pH regulation, we identified acidic and alkaline hits that encode for lysosome-resident proteins as defined by 3-fold enriched proteins identified previously through lysosome immunoprecipitation of lysosomes from iNeurons^42^ (**Extended Data Fig. 2c, Table S2**). As expected, the GO categories of this subset of hits were enriched for response to amino acid stimulus, regulation of mTOR signaling and lysosomal transport (**Extended Data Fig. 2d**). Notably, *PQLC2* (aka *SLC66A1* in humans and *laat-*1 in *C. elegans*) was the top hit among the genes encoding lysosomal components whose knockdown led to lysosomal alkalinization. In addition, it was recently found to be expressed at lower levels in aged, transdifferentiated neurons^43^, suggesting a possible role in aging and neurodegeneration. Thus, we further investigated the mechanism by which it regulates lysosomal pH.

PQLC2 mediates the efflux of basic amino acids (arginine, lysine and histidine) from the lysosomal lumen^44,45^, with loss of PQLC2 resulting in the luminal accumulation of basic amino acids ^46^. PQLC2 also recruits the C9Orf72-SMCR8-WDR41 complex to the lysosomal surface^47^. We generated both CRISPRi and CRISPR knock-out PQLC2 iPSC lines to validate the lysosomal pH phenotype (**Extended Data Fig. 3a,b**). To confirm the *PQLC2* screen phenotype, we transduced both *PQLC2* knockout and knockdown iPSCs with FIRE-pHLy. Upon neuronal differentiation, both of these *PQLC2* loss-of-function lines exhibited alkaline lysosomal pH by FIRE-pHLy ratio (**Fig. 3a,b** **and Extended Data Fig. 4a**) and dextran-based pH-measurement (**Fig. 3c** **and Extended Data Fig. 4b**). These findings demonstrate that *PQLC2* represents a robust hit whose normal function is required for lysosomal pH maintenance.

**Figure 3:**
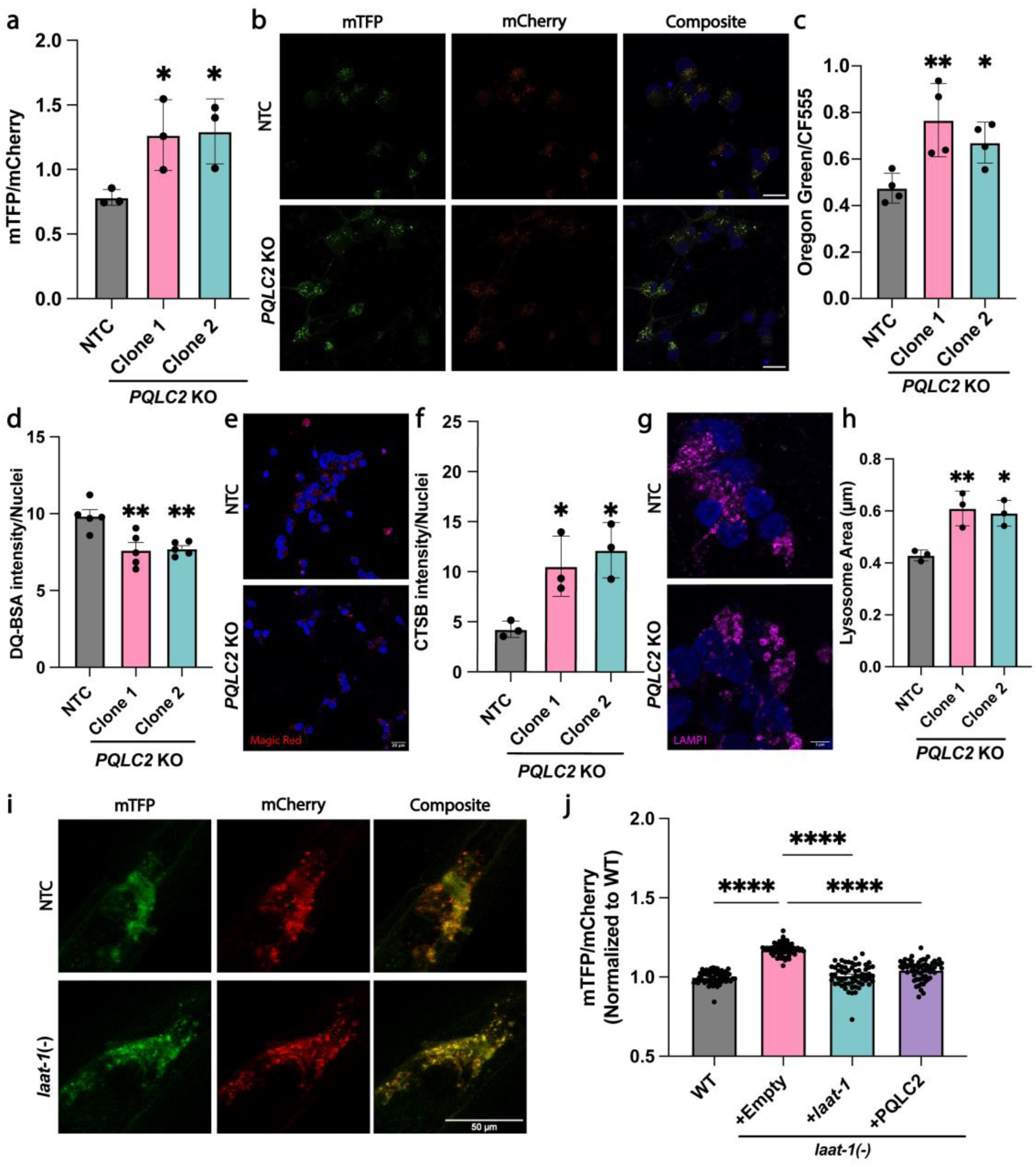
PQLC2 regulates neuronal lysosomal pH and lysosome function *in* vitro and *in vivo*. **a,** Day 14 PQLC2 knockout neurons have increased mTFP:mCherry ratio (n=3 independent experiments). **b,** Representative images of PQLC2 knockout iNeurons (scale bar = 20 μm)**. c,** Dextran-based lysosomal pH measurement in day 14 PQLC2 knockout neurons has increased Oregon green-488 to CF-555 ratio (n=4 independent experiments). **d,** DQ-BSA fluorescence intensity is decreased in day 14 PQLC2 knockout iNeurons compared to control cells**. e,** Representative image of Magic Red CTSB staining in PQLC2 knockout neurons (scale bar = 20 μm). **f,** PQLC2 knockout neurons demonstrated increased cathepsin B activity as measured by Magic Red CTSB. **g,** Representative image of LAMP1 staining in PQLC2 knockout neurons showing increased lysosome area (scale bar = 5 μm). **h,** Quantification of lysosomal area as marked by LAMP1 showing increased lysosomal area in PQLC2 knockout neurons (n=3 independent experiments). **i,** Representative images of *laat-1* mutant *C. elegans* expressing neuronal FIRE-pHLy. **j,** Neuronal lysosomal pH is increased in *C. elegans laat-1(-)* and is rescued by both *laat-1* or human *PQLC2* expression. * p <0.05, ** p <0.01, *** p <0.001, **** p <0.0001.

We next tested whether PQLC2 was the primary export pathway for cationic amino acids from the lysosome in iNeurons. We used NS560, a fluorescent compound that binds free amino acids, to assess amino acid levels in lysotracker-positive vesicles^48^. As expected, the amount of free amino acid was increased in PQLC2 knockout iNeurons (**Extended Data Fig. 4c**), which aligns with previous work in HEK293 cells showing cationic amino acid accumulation upon loss of PQLC2^46^.

Lysosomal pH can directly regulate lysosomal protease activity^12^, thus, we tested the degradative capacity of *PQLC2* KO cells. Interestingly, *PQLC2* KO cells demonstrated decreased proteolysis when measured by DQ-BSA (**Fig. 3d**). This overall loss of proteolytic activity is notable as certain proteases, such as CTSB, have a broad pH range of activity between pH 4 and 7^13,49^ and, in *PQLC2* KO cells CTSB activity was elevated (**Fig. 3e,f**). Loss of *PQLC2* did not impact CTSD activity (**Extended Data Fig. 4d**) but caused an increase in lysosome area (**Fig. 3g,h**). Thus, when PQLC2 is lost, lysosomal homeostasis appeared to be dysregulated with decreased degradative capacity and increased lysosome size.

Finally, we sought to establish the *in vivo* relevance of these findings. In *C. elegans, laat-1* is the PQLC2 homolog^44^. Loss of *laat-1* in *C. elegans* expressing neuronal FIRE-pHLy increased mTFP:mCherry ratio, demonstrating lysosomal alkalinization *in vivo* (**Fig. 3i,j**). To confirm that this effect was specific to *laat-1* loss-of-function, we performed rescue experiments. Re-expression of *laat-1* restored pH to wild-type levels (**Fig. 3j**). Notably, human *PQLC2* expression was just as effective as *laat-1* (**Fig. 3j**), providing functional evidence of a conserved role for these two proteins in lysosomal pH regulation and highlighting the *laat-1* mutant as an effective *in vivo* model for studying PQLC2 biology.

### mTORC1 signaling does not underlie the increased lysosomal pH phenotype in PQLC2 knockout

To understand how loss of *PQLC2* impairs lysosomal acidification, we measured transcriptomic differences between control and PQLC2 KO iNeurons. Genes involved in mTORC signaling were among the differentially expressed transcripts (**Extended Data Fig. 4e,f**). As mTORC signaling has previously been implicated in lysosomal pH changes^27^, we asked if downstream mTORC targets were affected by the loss of PQLC2. Loss of *PQLC2* did not alter the phosphorylation status of the multiple mTORC targets compared to controls (**Extended Data Fig. 4g-k**). Therefore, the alkaline lysosomal pH seen in PQLC2 knockout iNeurons is not likely due to inhibition of mTORC signaling.

### Canonical functions of PQLC2 do not regulate lysosomal pH

To further understand how *PQLC2* loss disrupts lysosomal pH in iNeurons, we performed a series of targeted mutations of amino acids that either are predicted to have a functional role and/or are highly conserved charged residues (**Fig. 4a,b**). Expression of wild-type (WT) PQLC2 under the weak constitutive promoter, PGK, was sufficient to rescue lysosomal pH to control levels in iNeurons (**Fig. 4c**). In response to cationic amino acid starvation, PQLC2 recruits the CSW complex (C9orf72, SMCR8 and WDR41) to the lysosome^47^. To test if binding of the CSW complex was required for lysosomal pH maintenance, we re-expressed PQLC2 F49A and PQLC2 W78A, two mutations shown to prevent or reduce the binding of the CSW complex to PQLC2 respectively^50^. Both mutants were capable of rescuing lysosomal pH in PQLC2 knockdown iNeurons (**Extended Data Fig. 5a),** suggesting the interaction between the CSW complex and PQLC2 was not required for lysosomal pH homeostasis. We then tested if expression of the three CSW-complex members was required to maintain lysosomal pH. Knockdown of *C9orf72, WDR41* or *SMCR8* did not impact lysosomal pH (**Fig. 4d**). Based on these data, the increased lysosomal pH of PQLC2 knockout is not due to disrupted interaction with the CSW complex.

**Figure 4:**
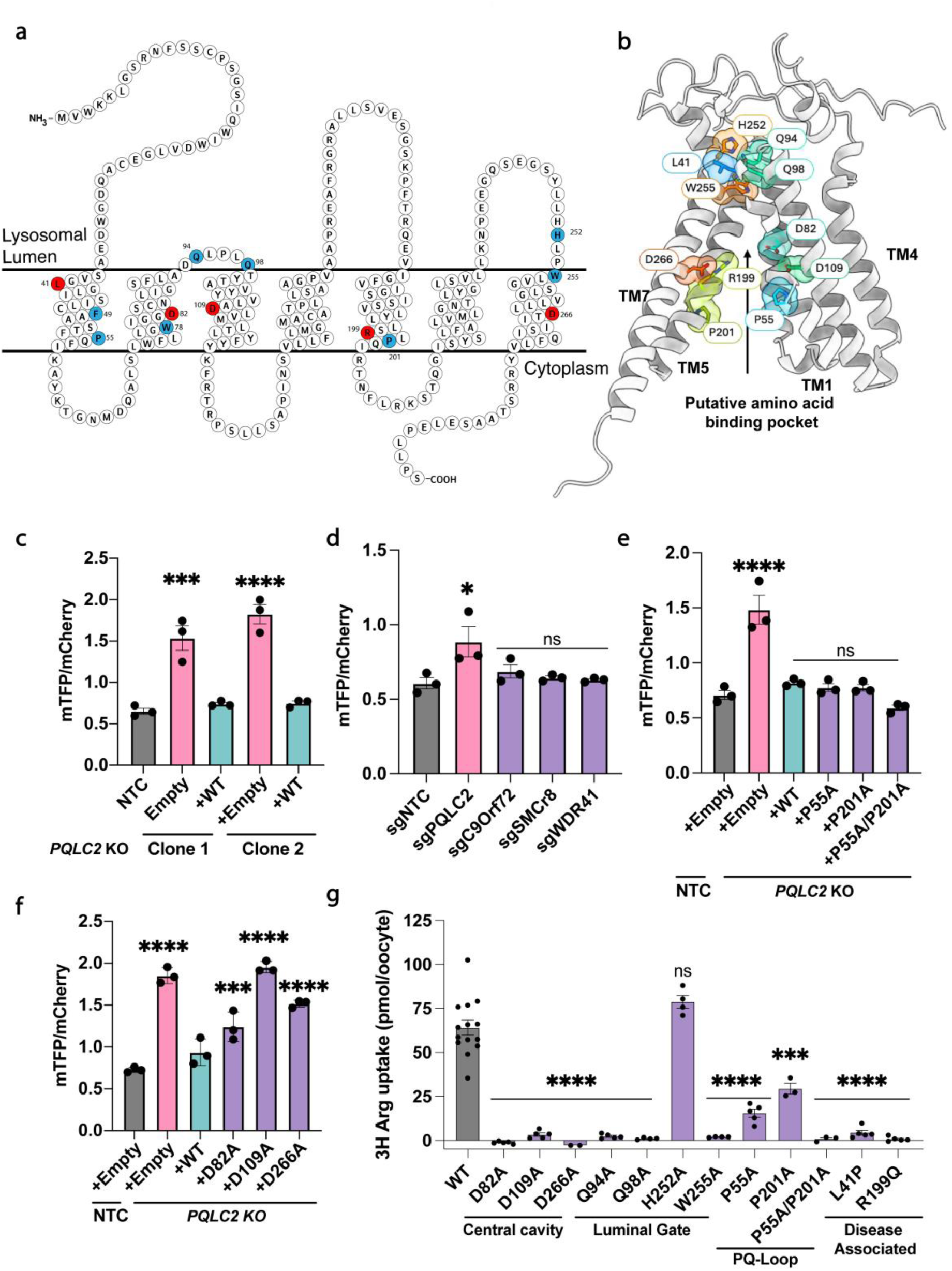
PQLC2 regulates lysosomal pH through a non-canonical function. **a,** Schematic of PQLC2 amino acids for essential lysosomal pH function. Red-labeled residues implicated in lysosomal pH regulation, while blue-labeled residues restored neuronal lysosomal pH. **b,** AlphaFold diagram of PQLC2 structure showing orientation of tested amino acid residues. **c,** Wild-type PQLC2 rescues lysosomal pH in day 14 neurons for two independent PQLC2 knockout clones (n= 3 independent experiments). **d,** Knockdown of PQLC2 or components of the C9orf72–SMCR8–WDR41 (CSW) complex by CRISPRi in day 14 iNeurons shows CSW complex is not required for lysosomal pH regulation (n= 3 independent experiments). **e,** Expression of PQ-loop mutants rescues lysosomal pH in day 14 PQLC2 knockout iNeurons (n= 3 independent experiments). **f,** Expression of aspartic acid mutants within the transmembrane domain of PQLC2 fail to rescue lysosomal pH in day 14 iNeurons (n= 3 independent experiments). **g,** PQLC2 alanine mutants in a *Xenopus* oocyte assay transporting 3H arginine, asterisk denotes comparison to wild-type PQLC2 (n=3-6 individual oocytes per mutation). *p < 0.05, **p < 0.01, or ***p<0.001, One-way ANOVA, asterisks denote comparison to nontargeting control unless otherwise noted.

The other canonical function of PQLC2 is the export of cationic amino acids from the lysosome. We tested PQLC2 PQ-loop mutants that abolish amino acid transport^44,51^. Somewhat surprisingly, PQ-loop mutants P55A, P201A and P55A/P201A each effectively restored normal lysosomal pH, demonstrating that the PQ-loop residues are not required for lysosomal pH acidification (**Fig. 4e**). These data uncouple amino acid export from lysosomal pH regulation, suggesting that PQLC2 maintains lysosomal pH through a mechanism distinct from its canonical transporter function.

We next targeted a series of amino acids in PQLC2—Q94, Q98, H252 and W255—that face the lysosomal lumen and are predicted to interact by AlphaFold (**Fig. 4b**). When expressed as alanine mutants in PQLC2 knockdown iNeurons, these luminal gate mutants still rescued lysosomal acidity (**Extended Data Fig. 5c**) indicating that these residues are also dispensable for pH regulation by PQLC2.

We then sought to identify residues within the core of PQLC2 that may play a role in lysosomal pH homeostasis. Three aspartic acids within the transmembrane central cavity of PQLC2, D82, D109 and D266, are predicted to be critical for amino acid transport. Expression of these central-cavity *PQLC2* mutants failed to rescue or only partially rescued lysosomal pH (**Fig. 4f**).

Finally, to comprehensively assess amino acid transport across this series of mutants, we measured arginine uptake following PQLC2 expression in *Xenopus* oocytes. As expected, the PQ-loop mutants exhibited markedly reduced arginine transport, retaining only ∼15–25% of wild-type activity (**Fig. 4g**). By contrast, mutation of the residues predicted to form the luminal gate of PQLC2 (Q94, Q98, and W255) completely abolished arginine transport, whereas mutation of H252 had no effect, identifying previously unrecognized residues that regulate PQLC2 transporter activity. In contrast, mutations within the central cavity failed to express, indicating that these residues are essential for correct transporter folding (**Fig. 4g** **and Extended Data Fig. 5d,e**). Together, these findings demonstrate that PQLC2-mediated export of basic amino acids can be functionally uncoupled from its role in regulating lysosomal acidification in iNeurons.

### Loss of PQLC2 increases tau levels *in vitro* and *in vivo*

We sought to determine if loss of *PQLC2* could impact clearance of tau, a protein that accumulates in Alzheimer’s disease and other neurodegenerative tauopathies. First, we measured phosphorylated tau (p-tau) in cultured iNeurons, as p-tau is more effectively cleared at a more acidic pH^52^. At post-differentiation days 21 and 42 (weeks 3 and 6), the relative level of p-tau was elevated in *PQLC2* KO iNeurons compared to controls (**Fig. 5a-c**). This increased p-tau suggests a potential defect in tau homeostasis upon the loss of PQLC2.

**Figure 5:**
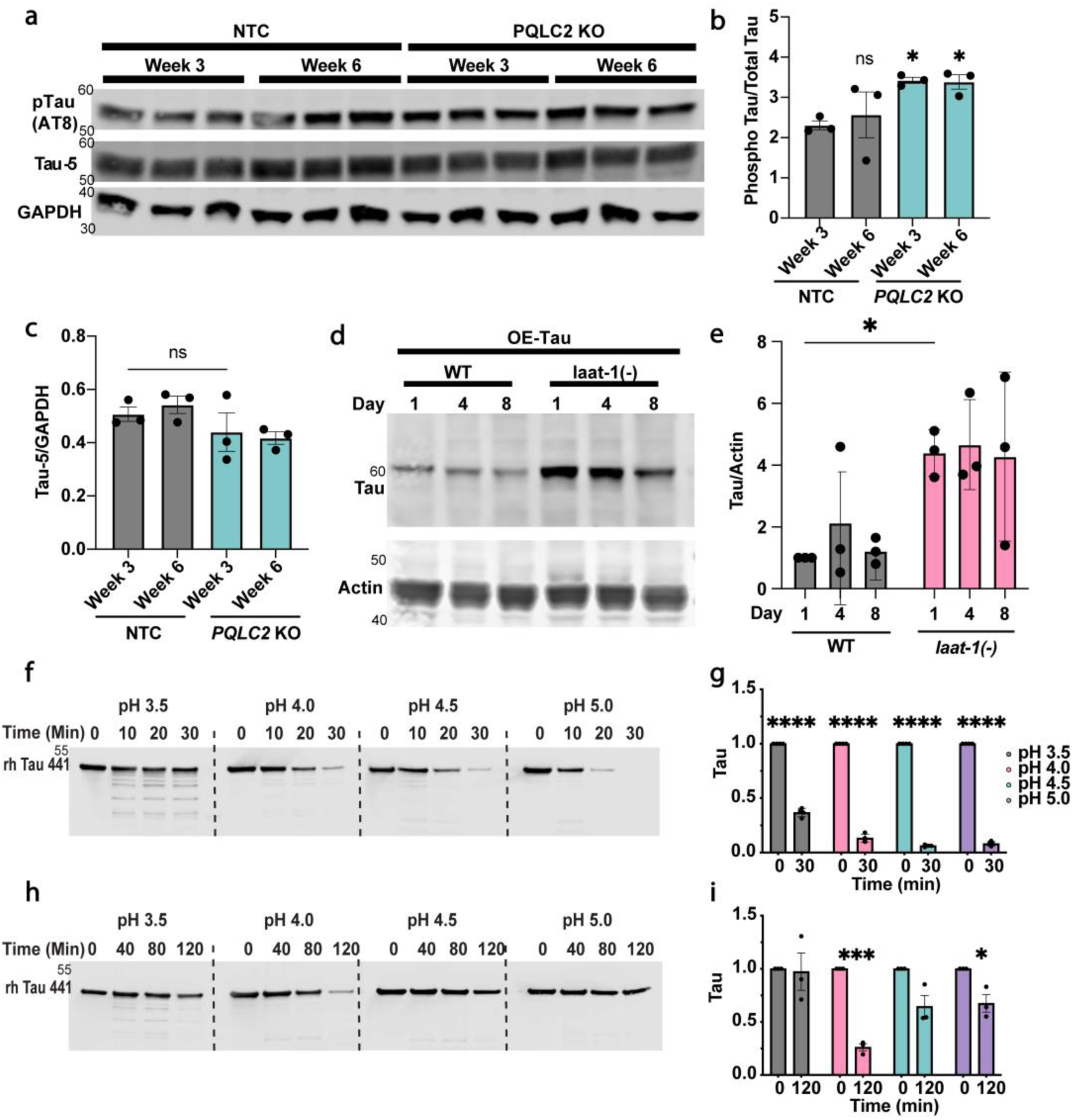
Loss of PQLC2 increases tau accumulation *in vitro* and *in vivo*. **a**, Western blot of phosphorylated tau (AT8) and total tau (Tau-5) protein levels in 3-week and 6-week iNeurons. **b**, Quantification of phosphorylated tau levels shown in **a**. finds increased phosphorylated tau levels at both 3- and 6- weeks in *PQLC2* knockout neurons compared with controls. Statistical significance was determined by one-way ANOVA with comparison to the 3-week non-targeting control (NTC). **c**, Quantification of total tau levels from the western blots shown in **a**, normalized to GAPDH, demonstrating that total tau levels are unchanged at both 3- and 6- weeks. Statistical significance was determined by one-way ANOVA with comparisons to the 3-week NTC. **d**, Western blot analysis of tau in Day 1, Day 4, and Day 8 control or *laat-1 mutant C. elegans* crossed with a human tau overexpression strain. **e**, Quantification of the western blots shown in **d**, demonstrating that *laat-1mutant* animals exhibit increased tau levels across early to mid-life. Two-way ANOVA identified genotype as a significant factor affecting total tau levels. **f**, Western blots of recombinant full-length human Tau-441 proteolysis by iPSC-derived lysosomes over 30 minutes *in vitro* at pH 3.5, 4.0, 4.5, and 5.0. **g**, Quantification of the 0- and 30-minute time points from the western blots shown in **f**. Statistical significance was determined by t-test. **h**, Western blots of recombinant full-length human Tau-441 proteolysis by iNeuron-derived lysosomes over 120 minutes *in vitro* at pH 3.5, 4.0, 4.5, and 5.0. **i**, Quantification of the 0- and 120-minute time points from the western blots shown in **h**. Statistical significance was determined by t-test. * p < 0.05, ** p < 0.01, *** p < 0.001.

To determine if loss of PQLC2 drove tau accumulation *in vivo,* we crossed the *C. elegans laat-1(-)* strain to a strain expressing human tau^53^. Animals mutant for *laat-1* exhibited increased tau accumulation during adulthood, spanning early to mid-adult ages (day 1, 4, and 8), compared to controls (**Fig. 5d,e).** These results suggest that *laat-1* expression and potentially lysosomal pH play a role in clearance of tau.

### Tau degradation by neuronal lysosomes requires a highly acidic pH

The efficiency by which lysosomes degrade specific substrates, including neurodegenerative disease proteins such as tau, can vary dramatically depending on pH^13,52,54^. Since neurons accumulate tau in Alzheimer’s disease and other tauopathies, we asked if iNeuron-derived lysosomes differ from lysosomes isolated from their parent iPSCs in their ability to proteolyze tau. We isolated lysosomes from iNeurons or iPSCs using the lyso-IP method^55^. When iPSC lysosomes were incubated with recombinant tau, they were able to efficiently degrade tau across a range of pH setpoints (**Fig. 5f,g**). In contrast, iNeuron-derived lysosomes degraded tau optimally at pH 4, with much lower efficiency at pH 4.5 and 5.0 (**Fig. 5h,i**). These results indicate that neuronal lysosomes have more acidic pH requirements for tau degradation compared to non-neuronal cells such as iPSCs.

### *PQLC2* is a novel tauopathy risk gene

Given the importance of *PQLC2* in lysosomal acidification, the accumulation of tau in *PQLC2* knockout models, and the restricted pH profile for tau degradation by iNeurons, we wondered if *PQLC2* was implicated in neurodegenerative tauopathies. We thus queried a whole-genome sequencing dataset from a cohort of 782 individuals comprised of subjects with neurodegenerative disease (n = 682) and unaffected controls (n = 100) for potentially pathogenic variants in *PQLC2*. The sequencing identified an individual with an atypical primary tauopathy bearing some resemblance to corticobasal degeneration (CBD, a form of frontotemporal lobar degeneration with tau pathology) who had a rare heterozygous variant in *PQLC2* resulting in a leucine to proline coding change at position 41 (**Fig. 6a**). Immunohistochemistry performed on sections from the middle frontal gyrus of this individual demonstrated enlarged and abnormal appearing lysosomes in neurons, some of which also showed p-tau aggregates (**Fig. 6b**). These features were most prominent in cortical layers 5 and 6 (**Fig. 6b,c**). These data reveal a striking phenotypic overlap between the *PQLC2* knockout iNeurons and this individual carrying a PQLC2 L41P mutation.

**Figure 6:**
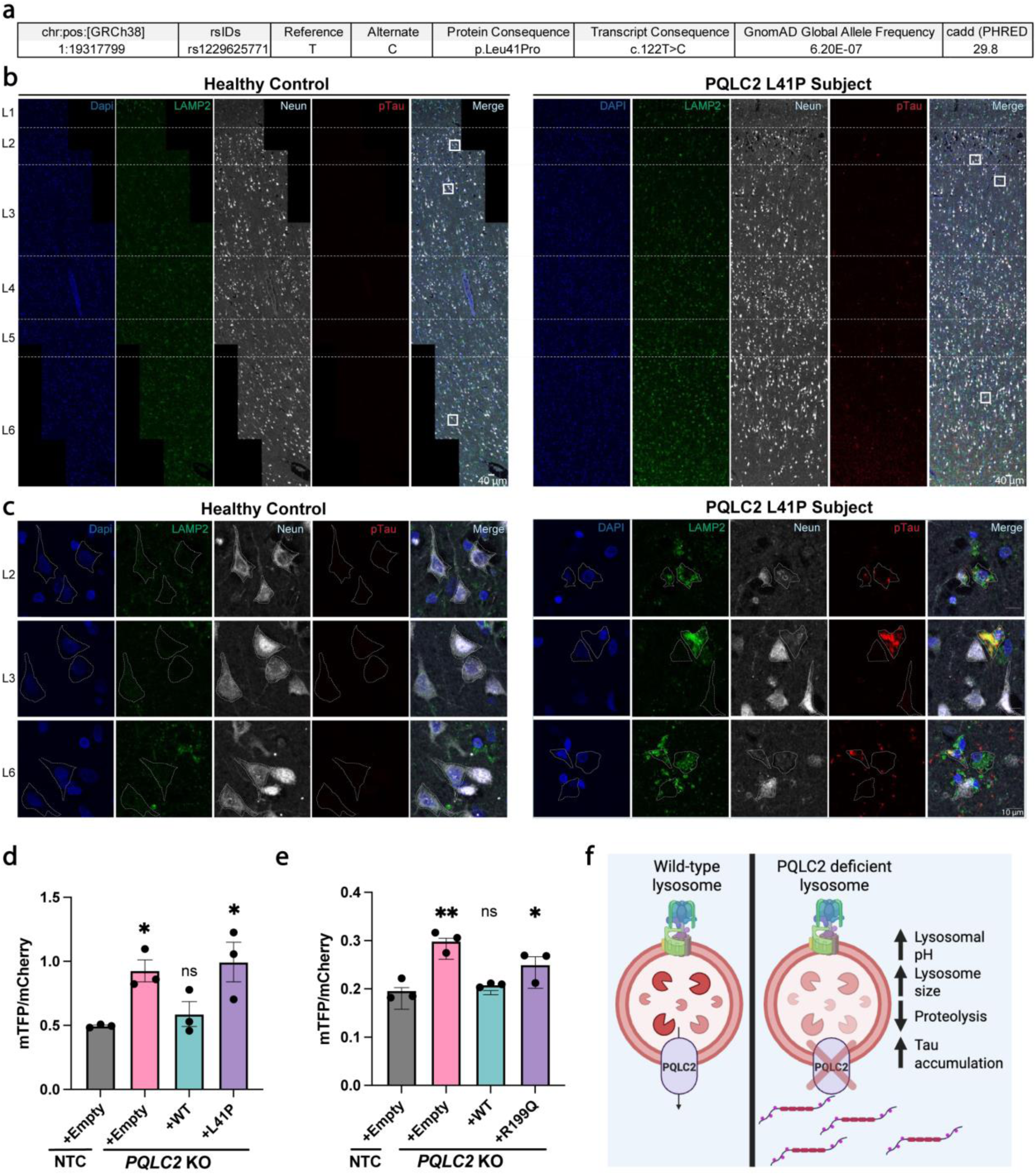
P*Q*LC2*/SLC66A1* is a novel tauopathy risk gene. **a,** A rare L41P missense variant (rs1229625771) was identified in an individual with pathology-confirmed tauopathy. The variant has a CADD score >25, predicting it is likely to be deleterious **b,** Tile scans of multiple layers of cortex from the middle frontal gyrus from an age-matched control or PQLC2 L41P subject stained for LAMP2, p-tau (CP13), and NeuN (scale bar = 40 μm) show prominent lysosome and p-tau changes in specific layers **c,** Representative insets shows more prominent LAMP2 and elevated p-tau in neurons from L2, L3 and L6 within the neurons of the L41P subject. **d,** Expression of tauopathy-associated PQLC2 L41P does not restore lysosomal pH in day 14 PQLC2 knockout neurons. **e,** Expression of the retinitis pigmentosa-associated PQLC2 R199Q mutation does not restore lysosomal pH in day 14 PQLC2 knockout neurons. **f,** Schematic showing how loss of PQLC2 affects neuronal health. * p < 0.05, or ** p < 0.01 One-way ANOVA, asterisks denote comparison to non-targeting control unless otherwise noted

To further investigate this disease-associated mutation, we tested the effect of the L41P variant on lysosomal pH. When expressed in PQLC2 knockout iNeurons, PQLC2 L41P failed to rescue lysosomal pH (**Fig. 6d)**. This suggests a potential mechanism by which this variant fails to clear tau, through alkalinizing lysosomal pH and disrupting lysosomal homeostasis and function. Previous reports have also linked homozygous PQLC2 R199Q to an age-associated form of retinitis pigmentosa, a rare degeneration of the retina^56^. When expressed in human PQLC2 knockout iNeurons, PQLC2 R199Q also failed to fully rescue lysosomal pH (**Fig. 6e**) highlighting the importance of lysosomal pH regulation by PQLC2 for neuronal health. Neither of these rare variants was competent to transport arginine (**Fig. 4g**) and had low levels of expression in the oocyte model (**Extended Data Fig. 5d,e**). The association of these two rare PQLC2 variants with neurodegeneration highlights the importance of lysosomal pH regulation in neurodegeneration and its key role in tau homeostasis (**Fig. 6f**).

## Discussion

Due to the association between proteostasis decline, lysosomal pH alkalinization and protein accumulation seen in aging and neurodegeneration^1,14,15^, understanding how lysosomal pH is regulated and how changes in pH affect substrate degradation are critical for understanding of neurodegenerative disease pathogenesis. The effects of individual genes related to neurodegenerative disease and aging on lysosomal pH have been studied in detail, but a comprehensive, unbiased approach towards understanding lysosomal pH had not been completed. A previous study using Lysosensor Yellow/Blue identified a small number of possible pH modifiers; however, due to their methodology, the screen primarily identified genes driving formation of hyper-acidic vacuoles in HAP1 cells^57^. We build on this approach using a more human relevant cell-type, iNeurons, and a robust, genetically encoded lysosomal pH sensor.

This study has identified genes not previously implicated in lysosomal pH regulation, including those important in mitochondrial translation, mitochondrial complex I assembly and respiration. While mitochondrial respiration has been known to impact lysosome biogenesis and function^58^, recent work has demonstrated lysosome-mitochondria contacts as a mechanism of lysosomal acidification^59,60^. Maintaining the lysosomal proton gradient is a dynamic and energetically costly process, requiring an estimated one ATP for the transport of three protons^61^. As mitochondria produce cellular ATP required for lysosomal acidification and lysosomal mitophagy is required for mitochondrial homeostasis, these findings support a mutual reliance between these two organelles. They also reconcile mitochondrial- and lysosome-centric theories of neurodegenerative disease. The question of which occurs first, mitochondrial or lysosomal failure, likely varies depending on individual cases. Nonetheless, maintaining lysosomal acidity and mitochondrial respiration during healthy aging could be complementary targets for preventing or delaying neurodegenerative processes. Genes whose knockdown leads to acidification, such as those of the UFMylation pathway, may represent processes whose loss lead to compensatory hyper-acidification. As UFMylation knockdown has been implicated in tau clearance^30,62,63^, though, it is likely that the compensation is not maintained with age.

PQLC2 is a strong lysosomal pH regulator that plays an additional role in tau homeostasis. This unexpected role for PQLC2 in tau clearance highlights the centrality of lysosomal pH in neurodegenerative disease pathogenesis, and that other lysosomal pH-regulating genes might represent previously unknown disease modifying targets. We have ruled out canonical functions of PQLC2 in lysosomal pH regulation and believe there is potential for a unique, previously unknown mechanism of lysosomal pH regulation involving PQLC2. This work highlights the diversity of proteins that affect neuronal lysosomal pH and emphasizes the role of lysosomal transporters beyond the V-ATPase in pH regulation. Further work is needed to dissect how many of these identified lysosomal pH modifiers work in concert to form a lysosomal pH regulatory network, and their potential association with disease.

Although this study was designed to be comprehensive, there are limitations. By performing the screen in iNeurons, there was a potential for strong pH modifiers to drop out by the day of analysis. Prior work has shown that predicted lysosomal pH modifiers, including V-ATPase components, have a survival phenotype unique to neuronal survival screens^64^, and could lead to an underestimation of a particular gene’s lysosomal pH phenotype. Additionally, despite the benefits that we have delineated, there are potentially pitfalls associated with using a genetically encoded sensor such as FIRE-pHLy due to its reliance on LAMP1 trafficking and potential for degradation. We attempted to address this with a secondary dextran-based counter screen, but both methods have the potential to misidentify or miss genuine lysosomal pH modifiers, and genes that were hits in one but not the other should not be discarded without further investigation. Further work will be needed to disentangle the differences in lysosomal pH measurement observed between the two methodologies.

## Acknowledgements

The authors would like to thank members of the Kao lab, Sarat Vatsavayai, Sevinc Jakab, Mor Alkaslasi and Avi Samelson for helpful discussions. We acknowledge the Gladstone Flow Cytometry Core, funded through NIH Grant P30 AI027763, and the UCSF Center for Advanced Technologies for sequencing supported by UCSF PBBR, RRP IMIA, and NIH 1S10OD028511-01 grants. We thank Shawn Ferguson for human PQLC2 cell lines.

## Methods

### Human iPSC Culture

Human iPSCs were cultured in Stemflex medium (ThermoFisher Scientific, #A3349401) on Matrigel (Corning, #354277) coated dishes. During passaging, cells were lifted using either ReLeSR (Stemcell Technologies, #100-0483) or accutase (Stemcell Technologies, #07922) and then replated in media supplemented with a 10 μM ROCK inhibitor (StemCell Technologies, #72304). Studies with human iPSCs at UCSF were approved by The Human Gamete, Embryo and Stem Cell Research (GESCR) Committee. Informed consent was obtained from the human subjects when the WTC11 or KOLF2.1J lines were originally derived^65,66^.

### Human iPSC-derived neuron culture

Differentiation of iPSCs into iNeurons was performed as previously described with minor modifications^67^. Briefly, iPSCs were plated onto Matrigel coated plates using N2-predifferentiation media consisting of Knockout DMEM (Thermo Fisher Scientific, #10-829-018) containing 2 ug/ml doxycycline (Clontech, # 631311), 1 ug/ml Rock inhibitor (StemCell Technologies, #72304), 1x N2 (Thermo Fisher Scientific, #17502048), 1x NEAA (Thermo Fisher Scientific, #11140050), 1 ug/ml NT-3 (Thermo Fisher Scientific, #450-03), 1 ug/ml BDNF (Thermo Fisher Scientific, #450-02) and 1 ug/ml laminin (Thermo Fisher Scientific, #23-017-015). The following day, media was exchanged for fresh media without rock inhibitor. After 72 hours of Ngn2 induction by doxycycline, iNeurons were replated using accutase onto plates or chamber slides (Ibidi, #80806) precoated with Polyethylenimine (PEI) (MilliporeSigma, #03880) and laminin. PEI-coated plates were prepared by diluting 50% PEI to 0.2% in 1x Borate buffer (ThermoFisher, #PI28341), added to cell culture plates and incubated at 37°C overnight. Plates were washed five times with water and allowed to dry completely prior to coating with 10 ug/ml laminin diluted in water.

After replating, iNeurons were maintained in N2/B27 media consisting of equal parts DMEM (Life Technologies Corporation, #11995073)/F12 (ThermoFisher, #11765062) and Neurobasal-A (Life Technologies Corporation, #10888022), 1x NEAA, 0.5x GlutaMAX (Thermo Fisher Scientific, #35050061), 0.5× N2 Supplement, 0.5x B27 supplement (Thermo Fisher Scientific, 17504044), 1 ug/ml NT-3 (Thermo Fisher Scientific, #450-03), 1 ug/ml BDNF (Thermo Fisher Scientific, #450-02) and 1 ug/ml laminin (Thermo Fisher Scientific, #23-017-015). Cells were subjected to a partial media change every three days to once a week depending on cell density.

### CRISPRi screening

For each genome-wide sub-library and each lysosomal pH sub-library screen, 45 million iPSCs in 3x T150s were infected with lentivirus at a multiplicity of infection (MOI) of ∼0.3 and after selected with 2 ug/ml of puromycin. Cells were selected for 5 days prior to differentiation. Differentiation of iPSCs was performed as previously described and plated on 10-cm PEI-coated dishes at a density of 10 million cells per plate. After two weeks of culture, cells were dissociated using the papain dissociation system (Worthington Biochemical Corporation, #9035-81-1). After dissociation, cells triturated 15 times with a 5-ml serological pipette and filtered through a 40µm cell strainer and pelleted at 200xg for 10 minutes. Cells were washed once in neuronal media, and resuspended in FACS buffer, 2% Fetal Bovine Serum, 1 mM EDTA, and 25 mM HEPES in Dulbecco’s Phosphate Buffered Saline (DBPS) (ThermoFisher, #14190144), and kept on ice until sorting. The top 30% and bottom 30% of mTFP:mCherry ratio was collected. After sorting, cells were pelleted at 200×g for 10 minutes, the supernatant was removed, and the pellet was frozen at −20°C. Genomic DNA was extracted with the NucleoSpin Blood S kit (Machery Nagel, # 740951.50). The sgRNA cassettes were amplified, pooled and sequenced as described^29^. Sequencing was analyzed as described for each sub-library^29^. Primary screens were analyzed using crispr_screen^68^. Briefly, raw sequencing reads were cropped and aligned to the protospacer sequences in our libraries using 2FAST2Q^69^. Aligned sgRNA counts were analyzed using an updated MAGeCK-iNC pipeline^29^ that uses a negative binomial distribution (https://noamteyssier.github.io/crispr_screen). Phenotypes and p-values for each gene were calculated and hit genes were called based on an FDR of 0.1. Hits were then combined, and gene set enrichment analysis was performed using ENRICHR^70–72^.

### pH sub-library cloning

A pool of sgRNA-containing oligonucleotides of top screen hits were synthesized by Twist Bioscience and cloned into an sgRNA expression vector as previously described^73^.

#### C. elegans Strains

The following Caenorhabditis elegans strains were used in this study:

AWK611, rocIs11[Prab-3::mTFP::LMP-1::mCherry];

CF3810, muIs216[Paex-3::huMAPT 4R1N + Pmyo-3::RFP];

AWK635, rocIs11[Prab-3::mTFP::LMP-1::mCherry + Pmyo-3::BFP]; laat-1(qx42) IV;

AWK653, laat-1(qx42) IV; muIs216[Paex-3::huMAPT 4R1N + Pmyo-3::RFP];

AWK672, laat-1(qx42) IV; rocIs11[Prab-3::mTFP::LMP-1::mCherry + Pmyo-3::BFP]; rocEx100[Peft-3::PQLC2::3×FLAG + Punc-122::GFP];

and AWK673, laat-1(qx42) IV; rocIs11[Prab-3::mTFP::LMP-1::mCherry + Pmyo-3::BFP];

rocEx101[Peft-3::laat-1(1)::3×FLAG + Punc-122::GFP].

The laat-1(qx42) allele^44^ was backcrossed four times to the N2E background before strain construction. Unless otherwise indicated, all strains were maintained at 20 °C under standard conditions as previously described^74^.

### Constructs, transgenes, and germ-line transformation

All plasmids were generated using the ClonExpress MultiS One Step Cloning Kit (Vazyme, Cat. No. C115). Transgenic animals were generated by microinjection into the syncytial gonad as previously described^75^. The extrachromosomal array rocEx100 (Peft-3::PQLC2::3×FLAG + Punc-122::GFP) and rocEx101 (Peft-3::laat-1(1)::3×FLAG + Punc-122::GFP) were generated by microinjection of pSS048 or pSS049, respectively, together with Punc-122::GFP as a co-injection marker into AWK635 (rocIs11[Prab-3::mTFP::LMP-1::mCherry + Pmyo-3::BFP]; laat-1(qx42)).

#### *C. elegans* confocal imaging and fluorescence quantification

Confocal imaging was performed on a Leica TCS SP8 microscope (Leica Microsystems) equipped with a 20× objective using LAS X software. L4 animals expressing mTFP::LMP-1::mCherry were immobilized in M9 buffer containing 5 mM levamisole (Thermo Fisher Scientific, #AC187870100) and mounted on 2% agarose pads (Precision Biosystems, #PBSA1705). mTFP and mCherry fluorescence were acquired sequentially using identical imaging settings for all samples within each experiment.

Fluorescence quantification was performed using Fiji (ImageJ, NIH). A lysosomal mask was generated from the mCherry channel by applying an intensity threshold of 20–255 and was subsequently applied to both the mTFP and mCherry channels to measure fluorescence intensity within lysosomal regions. Fluorescence intensities were normalized to the mean value of the wild-type control for each independent experiment before statistical analysis and data visualization.

### In vitro cleavage assays

Lysosomes were isolated from two confluent 15 cm dishes of iPSCs or iNeurons expressing Lyso-IP as previously described^55^, and eluted off the beads in PBS containing 0.1% NP-40. For in vitro cleavage assays, 1 µg of recombinant full-length 4N2R tau (rPeptide #T-1001–1) was incubated with isolated lysosomes from either iPSCs or iNeurons for the duration indicated at 37 °C in buffers ranging from pH 3.5 to 5.5 consisting of either 100 mM sodium citrate (pH 3.5) or 50 mM sodium acetate (pH 4.0, 4.5, 5.0 or 5.5). The reaction was quenched by moving the sample onto ice and adding NuPAGE 4X LDS (ThermoFisher #NP0007) and 10X reducing agent (Fisher #NP0009). Samples were then incubated for 10 min at 80°C. All samples were run on precast NOVEX 4–12% Bis–Tris gels (ThermoFisher #NP0321PK2) using MES buffer (ThermoFisher #NP0002) and transferred onto nitrocellulose membranes for Western blotting.

### Western blotting

*C. elegans* samples for western blotting were collected on days 1, 4 or 8 of adulthood. *C. elegans* pellets were lysed by repeated freeze–thaw cycles in a dry ice–ethanol bath and a 37 °C heat block, followed by sonication. Lysates were centrifuged at 13,000g for 5 min at 4 °C, and the supernatants were collected. Cell samples were lysed in 1x RIPA buffer with protease (ThermoFisher, #T3301) and phosphatase inhibitors (ThermoFisher, #4906845001). Protein concentrations were determined using the Pierce BCA Protein Assay Kit (ThermoFisher, #23225). Equal amounts of protein were mixed with NuPAGE LDS Sample Buffer and reducing agent and heated at 90°C for 10 min before electrophoresis. Samples were transferred onto nitrocellulose membranes. Membranes were blocked at room temperature with Odyssey Blocking Buffer (LI-COR #927-40,100) and incubated at 4 °C overnight with primary antibodies and 1 hour at room temperature with fluorescent secondary antibodies (LI-COR, #827-08364, #926-32210). Immunoreactive bands were visualized using a LI-COR Odyssey CLx image scanner and quantified with Image Studio version 5.5. Blots were quantified using ImageStudio and analyzed with GraphPad Prism (GraphPad Software, La Jolla, California USA).

### Antibodies

Antibodies used in this study are anti-ATP5A1 (Invitrogen, #459240), anti-LAMP2 (Biolegend, #354311), anti-EEA1 (BD Systems, #610457), anti-LAMP1 (Cell Signaling Technologies, #9091S) anti-p-Tau (AT8) (ThermoFisher, #MN1020), anti-p-Tau (CP13, gifted from Dr. Peter Davies) anti-LAMP2 (Abcam AB199946-1001), anti-NeuN (Synaptic Systems 266014), anti-Tau-5 (MilliporeSigma, #MAB361), anti-Tau (Abcam, ab76128), GAPDH (Abcam, #ab8245), and anti-β-actin (MilliporeSigma, MAB1501).

### iPSC cell line engineering

Genome editing of human KOLF2.1J was performed as described previously^29^. Briefly, the donor plasmid, CLYBL-TO-hNGN2-BSD-mApple (Addgene, #124229), was used to integrate a doxycycline-inducible human NGN2 transgene for neuronal differentiation into the CLYBL safe harbor locus. On the day of transfection, 2.5×10^5^ cells were plated per well in Stemflex containing 10μM Y-27632. Ribonucleoprotein (RNP) complexes of sgRNA and HiFi Cas9 nuclease were prepared and combined with 500 ng of donor DNA and 100 ng of pCE-mp53DD plasmid (Addgene, #41856). The mixture was delivered to cells using Lipofectamine™ Stem Transfection Reagent (ThermoFisher, #STEM00001). The day after transfection, cells were plated by serial dilution for clonal expansion. Antibiotic selection was applied 48-72 hours post-transfection, and mApple and BSD were removed with TAT-Cre Recombinase (MilliporeSigma, #SCR508).

Two monoclonal PQLC2 knockout iPSC lines were generated in the background of doxycycline inducible NGN2 KOLF2.1J iPSC^65^ using the CRISPR-Cas9 gene editing system. We designed a PQLC2 sgRNA targeting exon 3 (5’-AGCCTACAAGACGGGCAACA-3’) using CRISPOR^76^ and ligated into the plasmid PX459 as previously described^77^. Plasmids were transfected into 2.5×10^6^ iPSCs in a 6 well plate using Lipofectamine STEM. Just prior to transfection, media was replaced with 2 milliliters prewarmed of Opti-MEM (ThermoFisher, #31985070). After 4 hours, media was supplemented with 2 milliliters of Stemflex, and after 16 hours replaced with Stemflex media. After 48 hours, 2 μg/ml puromycin was added to select for transfected cells. Transfected cells were maintained in puromycin for 48 hours. Cells were diluted for clonal expansion and genotyping. DNA was collected from clones using the DNeasy Blood and Tissue kit (Qiagen, #69504), and gDNA was amplified using primers upstream and downstream of sgRNA target (5’- GTCCCCTATAAGGCGGCATC -3’, 5’- CAGGGTGAAGCAGGGTTAGG -3’). PCR product was sequenced for genotype information to confirm successful frameshift knockout of *PQLC2*.

### Fluorescence based lysosomal assays

#### For lysosomal pH measurement

Day 0 iNeurons were plated on eight-well chamber slides (Ibidi, 80806) coated with PEI/Laminin at a density of 2.5×10^5^ cells per chamber. For FIRE-pHLy measurement of lysosomal pH, FIRE-pHLy expressing cells were fixed on day 14 in 4% PFA. Fixed cells were washed twice with DPBS and imaged on an inverted confocal line-scanning microscope (DMi8 CS Bino, Leica Microsystems Inc., Wetzlar, Germany).

For dextran-conjugate pH measurement, 125 ug/ml Oregon Green-488 Dextran (Thermo Fischer Scientific, #D7170) and 20 µg/ml CF555-dextran (Biotium, #80112) were added to neuronal media and incubated overnight the day prior to imaging. The following morning, the cells were gently washed with fresh, warmed neuronal media five times and then the cells were incubated in fresh media for four hours. Dextran assays were imaged live, and media was exchanged to live cell imaging media which consisted of Hibernate-A low fluorescence media (ThermoFischer, NC0442869) supplemented with 0.5X N2, 1X B27, 1 µg/ml BDNF, and 1 µg/ml NT-3. Cells were maintained at 37°C using a humidified incubator chamber and imaged using an inverted confocal line-scanning microscope (DMi8 CS Bino, Leica Microsystems Inc) with either a 20x air objective, 40x oil-immersion objective or a 63x oil-immersion objective lens. Fluorescence images were acquired with sequential scanning between frames on the LAS X SP8 Control Software system using preset channel settings. Randomly imaged fields were processed (background subtraction, thresholding), and lysosomal pH ratios were calculated using a CellProfiler pipeline^78^. Briefly, puncta were segmented for pH insensitive fluorescence (mCherry or CF555) and used as a mask to measure both the pH sensitive and insensitive signal per puncta. The ratio for each punctum was taken and averaged to generate an average ratio per image.

#### For lysosomal substrate assays

Day 0 iNeurons were plated on eight-well chamber slides (Ibidi, 80806) coated with PEI/Laminin at a density of 2.5×10^5^ cells per chamber. On day 14 either: DQ BSA-Red (ThermoFisher, #D12051) was added at 50 µg/ml in neuronal media, and incubated for four hours, Pepstatin A, BODIPY™ FL Conjugate (ThermoScientific, P#12271) was added at 1 µM for one hour, NS560 (gifted from Dr. Timothy Glass) was added at 5µM for 30 minutes, lysotracker deep-red at 50 ng/ml (ThermoFisher, #L12492) or Magic Red Fluorescent Cathepsin B Assay Kit (Antibodies Inc, #938) was added at a 1:250 dilution in neuronal media for 30 minutes. After incubation, cells were gently washed five times with pre-warmed N2/B27 media and imaged in pre-warmed live cell imaging media.

#### Image analysis

Randomly imaged fields were processed (background subtraction, thresholding), and lysosomal pH ratios were calculated using a CellProfiler pipeline^78^. For lysosomal pH measurements, puncta were segmented for pH insensitive fluorescence (mCherry or CF555) and used as a mask to measure both the pH sensitive and insensitive signal per puncta. The ratio for each punctum was taken and averaged to generate an average ratio per image. For other lysosome assays, puncta were segmented, and integrated intensity was measured per puncta. These puncta were assigned to the nearest nuclei to derive per-cell averages per image to normalize for differences in cell density in different images.

### Immunocytochemistry and colocalization analysis

iNeurons were fixed on day 10 in 4% PFA for 15 minutes at room temperature, washed three times with DPBS, and permeabilized/blocked in blocking buffer (0.05% saponin, 5% normal goat serum in DPBS) for one hour. Cells were incubated with primary antibodies against either LAMP2, LAMP1, ATP5A1, or EEA1 overnight at 4°C, followed by fluorescent secondary antibodies for two hours at room temperature. Nuclei were counterstained with Hoechst 33342 for 10 minutes prior to imaging.

Colocalization between FIRE-pHLy (mCherry channel) and each marker was quantified using Pearson’s correlation coefficient with CellProfiler. Lysosome size was assessed by segmenting and measuring LAMP1 area in either wild-type or *PQLC2* knockout iNeurons in ImageJ/FIJI (v2.14.0/1.54f).

### RNA Sequencing and differential expression analysis

RNA sequencing was performed by culturing 1×10^6^ iNeurons on PEI/laminin coated plates. On day 14, cells were scraped in ice cold DPBS and pelleted at 500xg for 5 minutes. These pellets were then resuspended in DNA/RNA shield (Zymo Research, #R1100-50). Samples were submitted for Plasmidsaurus RNA-Seq (3′ end counting, Illumina NovaSeq X Plus). Gene-level quantification and differential gene expression analysis were performed using the Plasmidsaurus RNA-Seq analysis pipeline, using the human (GRCh38_114) Ensembl genome as reference.

### PQLC2 plasmid cloning

Wild-type PQLC2 was ordered codon-optimized from Twist Bioscience. This codon optimized gene was inserted into the backbone pLenti PGK Blast DEST (w524-1) which was a gift from Eric Campeau & Paul Kaufman (Addgene, #19065) using Gibson Assembly (New England Biolabs, #E5510S). PQLC2 mutants were generated by PCR to generate DNA fragments with our intended mutation and assembled using Gibson Assembly. cDNA was amplified using 1x Platinum Superfi II Mastermix (Thermo Fisher Scientific, #12368010) with 1x Evagreen dye (Biotium, #31000) at a 1x concentration and ROX Reference Dye 25 uM (Biotium, #29052) at 0.5 uM concentration. Primers used for *PQLC2* mRNA were (5’- gcctgggcttgatctccatt-3’, 5’- cacagccgtgtaggtctgc-3’), primers for *GAPDH* mRNA were (5’-ggtgtgaaccatgagaagtatga-3’, 5’-gagtccttccacgataccaaag-3’).

#### Expression and analysis in *Xenopus* oocytes

Human PQLC2 without its C-terminal dileucine-type sorting motif (PQLC2-LL/AA, hereafter referred to as WT) was subcloned with a C-terminal GFP into the pFAW vector for microscopy and transport assays. Variants of the protein were generated by site-directed mutagenesis and verified via sequencing. *In vitro* transcription of wild type (WT) and mutant *PQLC2* was carried out using AmpliCapTM-Max T7 high yield message maker kit (Cellscript). *Xenopus laevis* oocytes were defolliculated manually. They were then injected with 50 nl of nuclease-free water or 50 ng of human *PQLC2* WT or mutant mRNA and were incubated for 40-42 hr at 17 °C in Modified Barth’s storage solution at pH 7.4 (88 mM NaCl, 1 mM KCl, 1.68 mM MgSO_4_. 7H2O, 10 mM HEPES and 2.4 mM NaHCO_3_) supplemented with 0.05 mg/ml gentamicin.

Oocytes with high expression were selected under the epifluorescence microscope Leica Thunder. Computational clearing was applied using the THUNDER Large Volume Computational Clearing algorithm to remove out-of-focus background and improve image contrast using LAS X software.

Radiotracer flux analysis was performed in 96 mM NaCl, 2 mM KCl, 1 mM MgCl2, and 1.8 mM CaCl2 buffered with 5 mM MES adjusted to pH 5.0 with NaOH (ND96 buffer). Oocytes were incubated in ND96 with 2 μCi of [3H]arginine and 100 μM of non-radiolabeled arginine. Incubation time for uptake was fixed to 15 min. Uptake was stopped by two ice-cold ND96 washes at pH 7.5. Oocytes were lysed in 20 mM Tris pH 7.5 with 1% triton X-100, transferred into Ultima Gold (PerkinElmer) scintillation liquid and intracellular radioactivity was counted individually for each oocyte, using a Wallac scintillation counter.

### Identification of PQLC2 variant

Potentially pathogenic variants in PQLC2 were identified within a cohort of 782 individuals including neurodegenerative disease cases (n = 682) and unaffected controls (n = 100) recruited through the University of California, San Francisco (UCSF) Memory and Aging Center. Potentially pathogenic variants were defined by filtering for rare protein-coding variants using global population allele frequency data from gnomADv4.1.1 and deleteriousness predictions based on CADD v1.7 scores.

### Human brain tissue staining

Formalin-fixed, paraffin-embedded tissue sections were cut at 20 μm thickness from blocks representing the middle frontal gyrus. Sections were mounted on slides and dry-heated at 60°C for one hour. For deparaffinization, slides were washed three times in 100% Xylene for 5 minutes each, followed by 5 washes with 100% ethanol for 3.5 minutes, and a final wash with 95% ethanol for 3.5 minutes. We then performed antigen retrieval by autoclaving the sections for 5 minutes at 250°F in 0.01M Citrate buffer pH 6.0. After cooling, the slides were rinsed with 1x PBS containing 0.25% triton-X-100 for 5 minutes on shaker and then incubated in blocking buffer (10% goat serum, 3% bovine serum albumin, 0.25% triton-X-100 in 1x PBS) for one hour on shaker at room temperature. Primary antibodies [rabbit anti-Lamp2 (1:500), Abcam AB199946-1001; mouse anti-tau, phos Ser202 (CP13, 1:1000), gifted from Dr. Peter Davies; guinea pig anti-NeuN (1:100), Synaptic Systems 266014] were diluted in blocking buffer. The slides were incubated with primary antibodies overnight at room temperature. Next day, the slides were rinsed three times with 1x PBS containing 0.25% triton-X-100 for 5 minutes each on shaker. Secondary antibodies (Invitrogen) and DAPI were diluted in 1x PBS containing 0.25% triton-X-100 and added to the slides. The slides were incubated at room temperature for two hours. Slides were then rinsed with 1x PBS containing 0.25% triton-X-100 three times for 5 minutes each on shaker. For quenching autofluorescence, we treated the slides with 0.2% Sudan Black B dissolved in 70% ethanol. Briefly, slides were rinsed for 1 minute in deionized water, 5 minutes in 70% ethanol and then incubated for 20 minutes in 0.2% Sudan Black B at room temperature on shaker. Excess Sudan Black B was then removed by rinsing in 1x PBS containing 0.25% triton-X-100 for 3 minutes, with 1x PBS for 5 minutes and finally with deionized water for 5 minutes on shaker. The slides were mounted using Fluoromount G (Thermo Fisher, #00-4958-02) and coverslips were sealed with nail polish. Slides were imaged using Leica confocal SP8 using 63X oil immersion objective.

### Statistical Analysis

Details of the statistical test used for each experiment are described within each figure legend along with sample size and significance. For comparisons between two groups, unpaired two-tailed Student’s t-test was used, for comparisons among more than two groups, one-way ANOVA with Dunnett’s post-hoc correction was applied. For *C. elegans* tau accumulation data across multiple days of adulthood, two-way ANOVA with time and genotype as factors was performed. All data are represented as mean ± standard error. Statistical analysis was performed using GraphPad Prism 11 (GraphPad Software, La Jolla, California USA). For all tests, a p-value < 0.05 was considered significant.

## REFERENCES

1. Klaips, C. L., Jayaraj, G. G. & Hartl, F. U. Pathways of cellular proteostasis in aging and disease. J. Cell Biol. 217, 51–63 (2017).

2. Lim, C.-Y. & Zoncu, R. The lysosome as a command-and-control center for cellular metabolism. J. Cell Biol. 214, 653–664 (2016).

3. Zoncu, R. et al. mTORC1 Senses Lysosomal Amino Acids Through an Inside-Out Mechanism That Requires the Vacuolar H+-ATPase. Science 334, 678–683 (2011).

4. Cao, Q., Yang, Y., Zhong, X. Z. & Dong, X.-P. The lysosomal Ca2+ release channel TRPML1 regulates lysosome size by activating calmodulin. J. Biol. Chem. 292, 8424–8435 (2017).

5. Settembre, C., Fraldi, A., Medina, D. L. & Ballabio, A. Signals from the lysosome: a control centre for cellular clearance and energy metabolism. Nat. Rev. Mol. Cell Biol. 14, 283–296 (2013).

6. Srivastava, J., Barber, D. L. & Jacobson, M. P. Intracellular pH Sensors: Design Principles and Functional Significance. Physiology 22, 30–39 (2007).

7. Kane, P. M. Disassembly and reassembly of the yeast vacuolar H(+)-ATPase in vivo. J. Biol. Chem. 270, 17025–17032 (1995).

8. Forgac, M. Vacuolar ATPases: rotary proton pumps in physiology and pathophysiology. Nat. Rev. Mol. Cell Biol. 8, 917–929 (2007).

9. Dong, X. et al. PI(3,5)P2 controls membrane trafficking by direct activation of mucolipin Ca2+ release channels in the endolysosome. Nat. Commun. 1, 38 (2010).

10. Hu, M. et al. Parkinson’s disease-risk protein TMEM175 is a proton-activated proton channel in lysosomes. Cell 185, 2292–2308.e20 (2022).

11. Graves, A. R., Curran, P. K., Smith, C. L. & Mindell, J. A. The Cl-/H+ antiporter ClC-7 is the primary chloride permeation pathway in lysosomes. Nature 453, 788–792 (2008).

12. Turk, B. et al. Acidic pH as a physiological regulator of human cathepsin L activity. Eur. J. Biochem. 259, 926–932 (1999).

13. Sampognaro, P. J. et al. Mutations in α-synuclein, TDP-43 and tau prolong protein half-life through diminished degradation by lysosomal proteases. Mol. Neurodegener. 18, 29 (2023).

14. Hughes, A. L. & Gottschling, D. E. An early age increase in vacuolar pH limits mitochondrial function and lifespan in yeast. Nature 492, 261–265 (2012).

15. Sun, Y. et al. Lysosome activity is modulated by multiple longevity pathways and is important for lifespan extension in C. elegans. eLife 9, e55745 (2020).

16. Wallings, R., Connor-Robson, N. & Wade-Martins, R. LRRK2 interacts with the vacuolar-type H+-ATPase pump a1 subunit to regulate lysosomal function. Hum. Mol. Genet. 28, 2696–2710 (2019).

17. Lee, J.-H. et al. Lysosomal proteolysis and autophagy require presenilin 1 and are disrupted by Alzheimer-related PS1 mutations. Cell 141, 1146–1158 (2010).

18. Logan, T. et al. Rescue of a lysosomal storage disorder caused by Grn loss of function with a brain penetrant progranulin biologic. Cell 184, 4651–4668.e25 (2021).

19. Dehay, B. et al. Loss of P-type ATPase ATP13A2/PARK9 function induces general lysosomal deficiency and leads to Parkinson disease neurodegeneration. Proc. Natl. Acad. Sci. 109, 9611–9616 (2012).

20. Lee, J.-H. et al. Faulty autolysosome acidification in Alzheimer’s disease mouse models induces autophagic build-up of Aβ in neurons, yielding senile plaques. Nat. Neurosci. 25, 688–701 (2022).

21. Arotcarena, M.-L. et al. Acidic nanoparticles protect against α-synuclein-induced neurodegeneration through the restoration of lysosomal function. Aging Cell 21, e13584 (2022).

22. Zhou, N. et al. SLC7A11 is an unconventional H+ transporter in lysosomes. Cell 188, 3441–3458.e25 (2025).

23. Guo, J. et al. Aging and aging-related diseases: from molecular mechanisms to interventions and treatments. Signal Transduct. Target. Ther. 7, 391 (2022).

24. Canton, J. & Grinstein, S. Chapter 5 - Measuring lysosomal pH by fluorescence microscopy. in Methods in Cell Biology (eds Platt, F. & Platt, N.) vol. 126 85–99 (Academic Press, 2015).

25. Wolfe, D. M. et al. Autophagy failure in Alzheimer’s disease and the role of defective lysosomal acidification. Eur. J. Neurosci. 37, 1949–1961 (2013).

26. Chin, M. Y. et al. Genetically Encoded, pH-Sensitive mTFP1 Biosensor for Probing Lysosomal pH. ACS Sens. 6, 2168–2180 (2021).

27. Chin, M. Y. et al. Phenotypic Screening Using High-Content Imaging to Identify Lysosomal pH Modulators in a Neuronal Cell Model. ACS Chem. Neurosci. 13, 1505–1516 (2022).

28. Villa, S. et al. BiDAC-dependent degradation of plasma membrane proteins by the endolysosomal system. Nat. Commun. 16, 4345 (2025).

29. Tian, R. et al. CRISPR Interference-Based Platform for Multimodal Genetic Screens in Human iPSC-Derived Neurons. Neuron 104, 239–255.e12 (2019).

30. Samelson, A. J. et al. CRISPR screens in iPSC-derived neurons reveal principles of tau proteostasis. Cell 189, 1517–1534.e19 (2026).

31. Parra Bravo, C., et al. Human iPSC 4R tauopathy model uncovers modifiers of tau propagation. Cell 187, 2446–2464.e22 (2024).

32. Cao, Y. et al. DDRGK1, a crucial player of ufmylation system, is indispensable for autophagic degradation by regulating lysosomal function. Cell Death Dis. 12, 416 (2021).

33. Ratto, E. et al. Direct control of lysosomal catabolic activity by mTORC1 through regulation of V-ATPase assembly. Nat. Commun. 13, 4848 (2022).

34. Klein, Z. A. et al. Loss of TMEM106B Ameliorates Lysosomal and Frontotemporal Dementia-Related Phenotypes in Progranulin-Deficient Mice. Neuron 95, 281–296.e6 (2017).

35. Soyombo, A. A. et al. TRP-ML1 regulates lysosomal pH and acidic lysosomal lipid hydrolytic activity. J. Biol. Chem. 281, 7294–7301 (2006).

36. Leray, X. et al. Tonic inhibition of the chloride/proton antiporter ClC-7 by PI(3,5)P2 is crucial for lysosomal pH maintenance. eLife 11, e74136 (2022).

37. Bennett, N. K. et al. Defining the ATPome reveals cross-optimization of metabolic pathways. Nat. Commun. 11, 4319 (2020).

38. Ma, X. et al. Insights into the distinct membrane targeting mechanisms of WDR91 family proteins. Structure 32, 2287–2300.e4 (2024).

39. Qian, Y. et al. Functions of the α, β, and γ Subunits of UDP-GlcNAc:Lysosomal Enzyme N-Acetylglucosamine-1-phosphotransferase *. J. Biol. Chem. 285, 3360–3370 (2010).

40. Hutagalung, A. H. & Novick, P. J. Role of Rab GTPases in Membrane Traffic and Cell Physiology. Physiol. Rev. 91, 119–149 (2011).

41. Liv, N., Fermie, J., Ten Brink, C. B. M., de Heus, C. & Klumperman, J. Functional characterization of endo-lysosomal compartments by correlative live-cell and volume electron microscopy. Methods Cell Biol. 177, 301–326 (2023).

42. Hasan, S. et al. Multi-modal proteomic characterization of lysosomal function and proteostasis in progranulin-deficient neurons. Mol. Neurodegener. 18, 87 (2023).

43. Chou, C.-C. et al. Proteostasis and lysosomal repair deficits in transdifferentiated neurons of Alzheimer’s disease. Nat. Cell Biol. 27, 619–632 (2025).

44. Liu, B., Du, H., Rutkowski, R., Gartner, A. & Wang, X. LAAT-1 is the lysosomal lysine/arginine transporter that maintains amino acid homeostasis. Science 337, 351–354 (2012).

45. Jézégou, A. et al. Heptahelical protein PQLC2 is a lysosomal cationic amino acid exporter underlying the action of cysteamine in cystinosis therapy. Proc. Natl. Acad. Sci. 109, E3434–E3443 (2012).

46. Zhu, H. et al. Metabolomic profiling of single enlarged lysosomes. Nat. Methods 18, 788–798 (2021).

47. Amick, J., Tharkeshwar, A. K., Talaia, G. & Ferguson, S. M. PQLC2 recruits the C9orf72 complex to lysosomes in response to cationic amino acid starvation. J. Cell Biol. 219, e201906076 (2020).

48. Smith, M. R. et al. A Turn-On Fluorescent Amino Acid Sensor Reveals Chloroquine’s Effect on Cellular Amino Acids via Inhibiting Cathepsin L. ACS Cent. Sci. 9, 980–991 (2023).

49. Almeida, P. C. et al. Cathepsin B Activity Regulation: HEPARIN-LIKE GLYCOSAMINOGLYCANS PROTECT HUMAN CATHEPSIN B FROM ALKALINE pH-INDUCED INACTIVATION *. J. Biol. Chem. 276, 944–951 (2001).

50. Talaia, G., Amick, J. & Ferguson, S. M. Receptor-like role for PQLC2 amino acid transporter in the lysosomal sensing of cationic amino acids. Proc. Natl. Acad. Sci. U. S. A. 118, e2014941118 (2021).

51. Kawano-Kawada, M. et al. A PQ-loop protein Ypq2 is involved in the exchange of arginine and histidine across the vacuolar membrane of Saccharomyces cerevisiae. Sci. Rep. 9, 15018 (2019).

52. Lane-Donovan, C. et al. Tau phosphorylation at Alzheimer’s disease biomarker sites impairs its cleavage by lysosomal proteases. Alzheimers Dement. J. Alzheimers Assoc. 21, e70320 (2025).

53. Butler, V. J. et al. Tau/MAPT disease-associated variant A152T alters tau function and toxicity via impaired retrograde axonal transport. Hum. Mol. Genet. 28, 1498–1514 (2019).

54. Banay-Schwartz, M. et al. The pH dependence of breakdown of various purified brain proteins by cathepsin D preparations. Neurochem. Int. 7, 607–614 (1985).

55. Abu-Remaileh, M. et al. Lysosomal metabolomics reveals V-ATPase and mTOR-dependent regulation of amino acid efflux from lysosomes. Science 358, 807–813 (2017).

56. Millo, T. et al. Identification of autosomal recessive novel genes and retinal phenotypes in members of the solute carrier (SLC) superfamily. Genet. Med. 24, 1523–1535 (2022).

57. Lenk, G. M. et al. CRISPR knockout screen implicates three genes in lysosome function. Sci. Rep. 9, 9609 (2019).

58. Baixauli, F. et al. Mitochondrial Respiration Controls Lysosomal Function during Inflammatory T Cell Responses. Cell Metab. 22, 485–498 (2015).

59. Tian, Z. et al. Mitochondria acidify lysosomes through membrane contacts. Cell Rep. 45, (2026).

60. Liu, Q. et al. Mitochondria-lysosome coupling contributes to lysosome acidification and aging. Mol. Cell 86, 2425–2442.e10 (2026).

61. Freeman, S. A. & Grinstein, S. Is the Parkinson’s-associated protein TMEM175 a proton channel: Yay or nay? J. Cell Biol. 225, e202511084 (2025).

62. C, P. B., et al. Human iPSC 4R tauopathy model uncovers modifiers of tau propagation. Cell 187, (2024).

63. Yan, T. et al. The UFMylation pathway is impaired in Alzheimer’s disease. Mol. Neurodegener. 19, 97 (2024).

64. Tian, R. et al. Genome-wide CRISPRi/a screens in human neurons link lysosomal failure to ferroptosis. Nat. Neurosci. 24, 1020–1034 (2021).

65. Pantazis, C. B. et al. A reference human induced pluripotent stem cell line for large-scale collaborative studies. Cell Stem Cell 29, 1685–1702.e22 (2022).

66. Kreitzer, F. R. et al. A robust method to derive functional neural crest cells from human pluripotent stem cells. Am. J. Stem Cells 2, 119–131 (2013).

67. Fernandopulle, M. S. et al. Transcription Factor-Mediated Differentiation of Human iPSCs into Neurons. Curr. Protoc. Cell Biol. 79, e51 (2018).

68. Teter, O. M. et al. CRISPRi-based screen of autism spectrum disorder risk genes in microglia uncovers roles of ADNP in microglia endocytosis and synaptic pruning. Mol. Psychiatry 30, 4176–4193 (2025).

69. Bravo, A. M., Typas, A. & Veening, J.-W. 2FAST2Q: a general-purpose sequence search and counting program for FASTQ files. PeerJ 10, e14041 (2022).

70. Chen, E. Y. et al. Enrichr: interactive and collaborative HTML5 gene list enrichment analysis tool. BMC Bioinformatics 14, 128 (2013).

71. Kuleshov, M. V. et al. Enrichr: a comprehensive gene set enrichment analysis web server 2016 update. Nucleic Acids Res. 44, W90–97 (2016).

72. Xie, Z. et al. Gene Set Knowledge Discovery with Enrichr. Curr. Protoc. 1, e90 (2021).

73. Horlbeck, M. A. et al. Compact and highly active next-generation libraries for CRISPR-mediated gene repression and activation. eLife 5, (2016).

74. Brenner, S. The genetics of Caenorhabditis elegans. Genetics 77, 71–94 (1974).

75. Mello, C. C., Kramer, J. M., Stinchcomb, D. & Ambros, V. Efficient gene transfer in C.elegans: extrachromosomal maintenance and integration of transforming sequences. EMBO J. 10, 3959–3970 (1991).

76. Concordet, J.-P. & Haeussler, M. CRISPOR: intuitive guide selection for CRISPR/Cas9 genome editing experiments and screens. Nucleic Acids Res. 46, W242–W245 (2018).

77. Ran, F. A. et al. Genome engineering using the CRISPR-Cas9 system. Nat. Protoc. 8, 2281–2308 (2013).

78. Stirling, D. R. et al. CellProfiler 4: improvements in speed, utility and usability. BMC Bioinformatics 22, 433 (2021).

79. Alquezar, C. et al. TSC1 loss increases risk for tauopathy by inducing tau acetylation and preventing tau clearance via chaperone-mediated autophagy. Sci. Adv. 7, eabg3897 (2021).

