## Extended Data Fig. 1 for "A neuronal CRISPRi screen identifies PQLC2 as a lysosomal pH regulator controlling tau homeostasis"

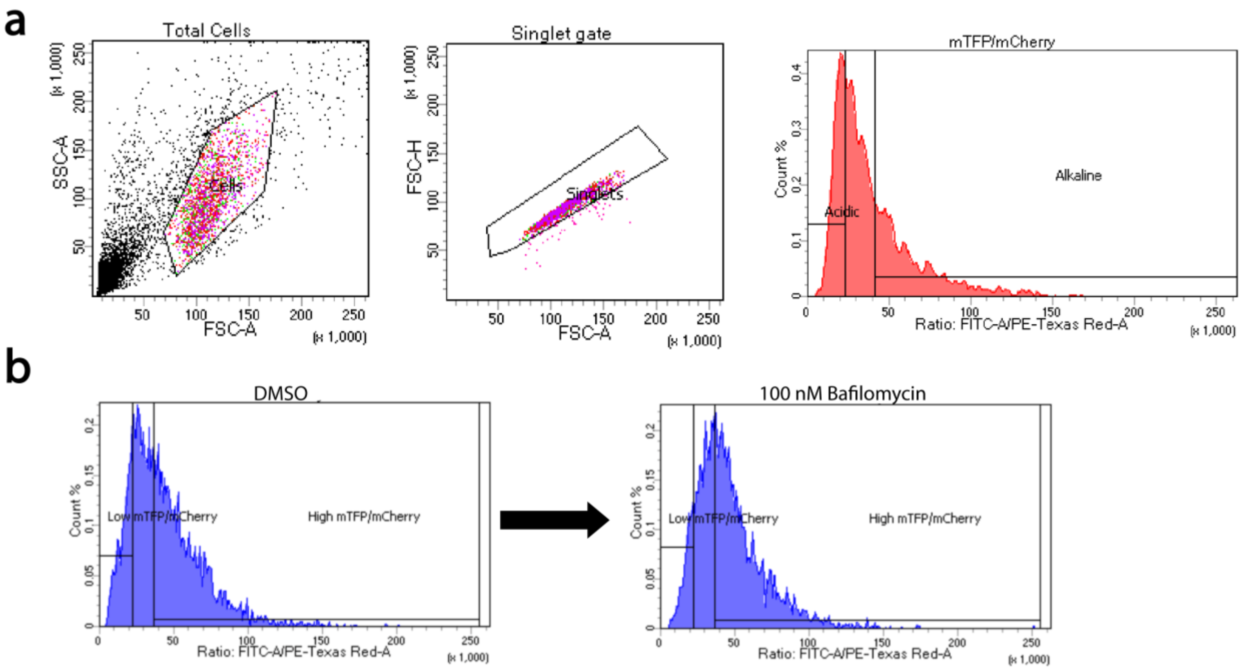


**Extended Data Figure 1: FIRE-pHLy is responsiv e to lysosomal pH changes measured by FACS sorting. a**, Representative gating strategy for sorting high mTFP:mCherry and low mTFP:mCherry populations of cells. **b**, Representative image showing increased mTFP:mCherry ratio in iNeurons as measured by FACS after 100 nM bafilomycin-A1 treatment overnight.
