## Extended Data Fig. 2 for "A neuronal CRISPRi screen identifies PQLC2 as a lysosomal pH regulator controlling tau homeostasis"

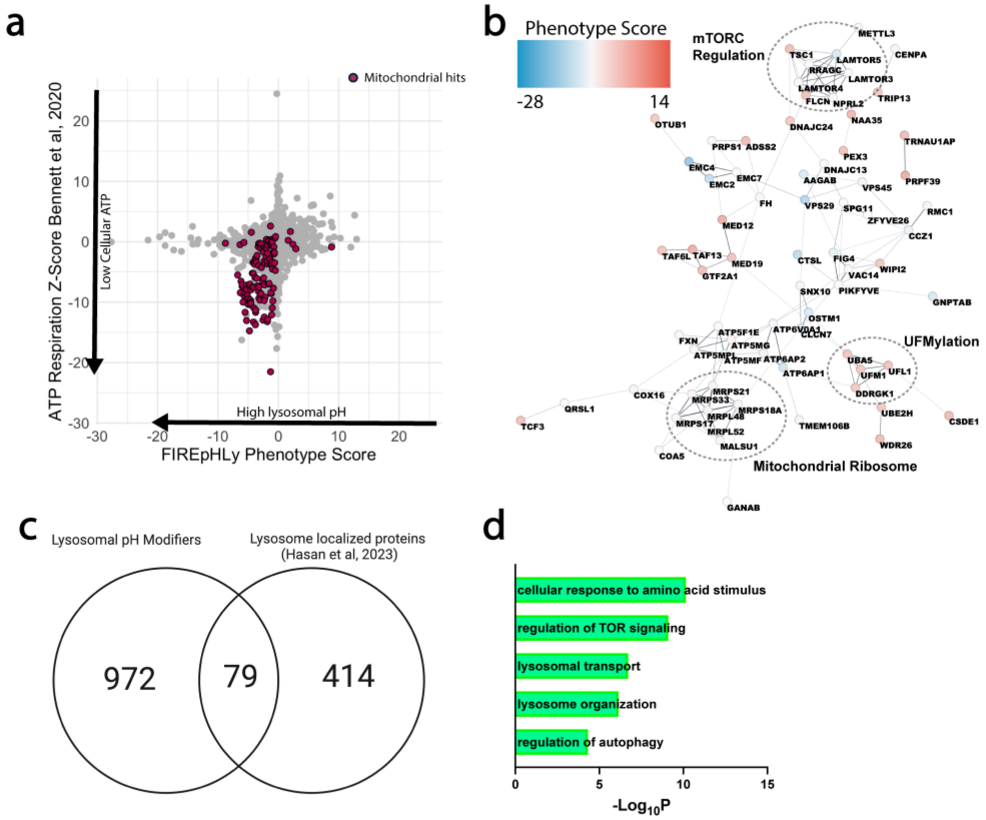


**Extended Data Figure 2: Comparisons of FIRE-pHLy CRISPRi screen results. a**, Comparison of FIRE-pHLy lysosomal pH screen phenotype scores against ATP-modifying hits under respiratory conditions reported by Bennett et al. 2020 highlighting mitochondrial lysosomal pH-modifying hits as reducing cellular ATP levels. **b**, STRING functional network of top lysosomal pH hits with singletons and doublets removed. **c,** Overlap between lysosomal pH-modifying hits and neuronal lysosome localized proteins reported by Hasan et al, 2023, as defined by threefold enrichment over cytosolic fraction. **d,** GO term enrichment of lysosome-localized lysosomal pH modifiers highlights mTORC regulation and lysosomal transporters as enriched biological processes.
