## Extended Data Fig. 3 for "A neuronal CRISPRi screen identifies PQLC2 as a lysosomal pH regulator controlling tau homeostasis"

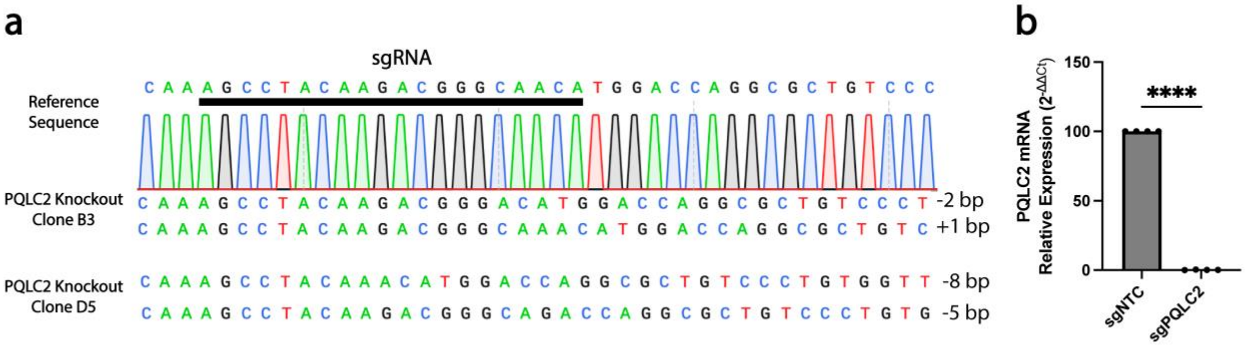


**Extended Data Figure 3: Validation of *PQLC2* iPSC lines. a,** Sequencing of two clones confirming frameshift mutations in both alleles of *PQLC2*. **b,** Relative mRNA levels of *PQLC2* in CRISPRi-mediated *PQLC2* knockdown neurons normalized to *GAPDH*. **** $p < 0.0001$.
