## Extended Data Fig. 4 for "A neuronal CRISPRi screen identifies PQLC2 as a lysosomal pH regulator controlling tau homeostasis"

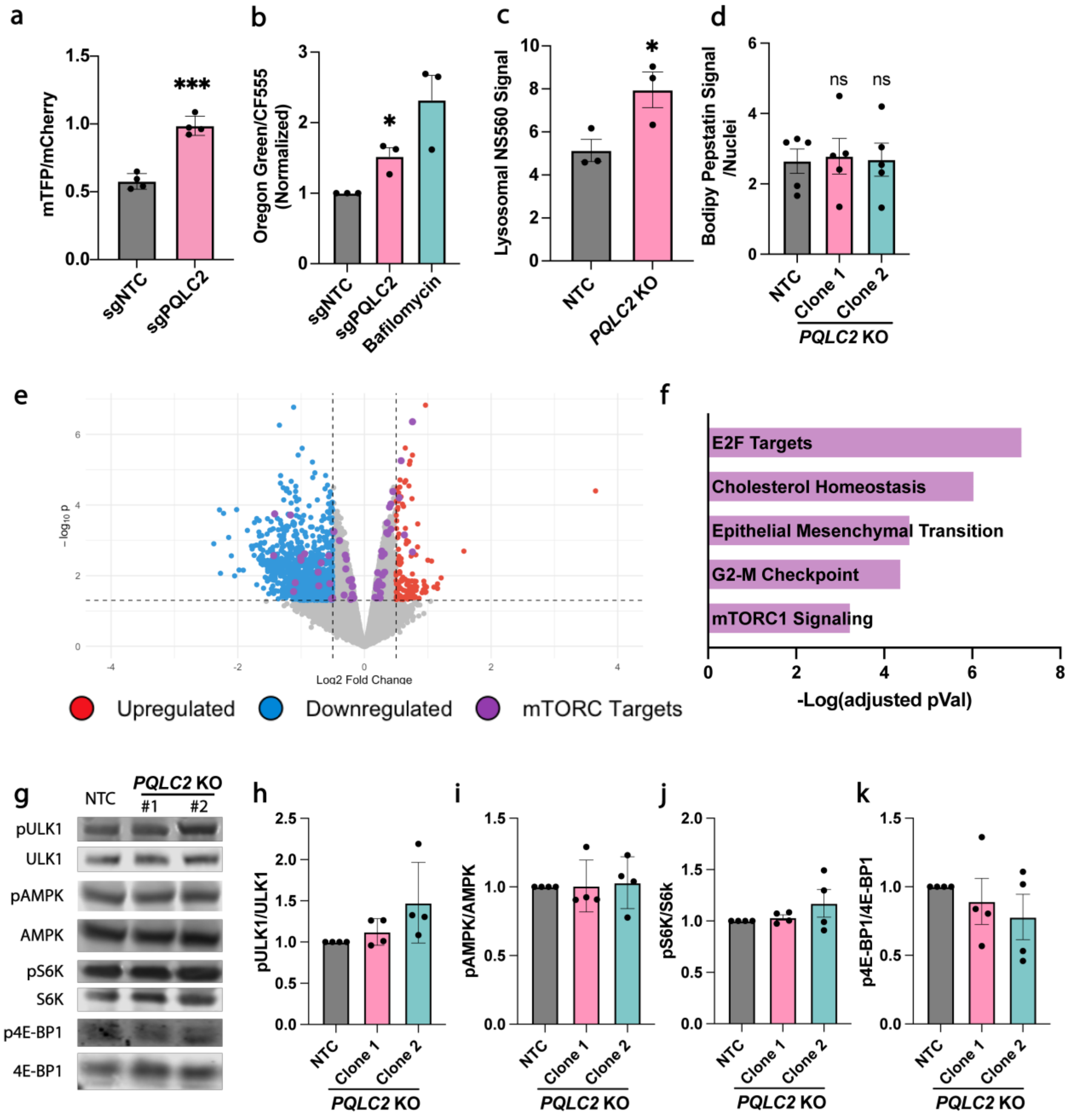


**Extended Data Figure 4: Loss of *PQLC2* does not disrupt neuronal mTORC1 signaling. a,** Day 14 *PQLC2* knockdown CRISPRi neurons have increased mTFP:mCherry ratio compared to non-targeting controls. **b**, Oregon green to CF555 ratio in day 14 *PQLC2* knockdown CRISPRi neurons show more alkaline lysosomal pH compared to controls. **c,** Lysosomal amino acid content measured by NS560 fluorescence in lysotracker positive puncta shows increased amino acid content in *PQLC2* knockout lysosomes. **d,** Bodipy-pepstatin fluorescence normalized to nuclei count demonstrates unchanged levels of active CTSD upon PQLC2 knockout. **e,** Volcano plot of differentially expressed genes (DEGs) in day 14 *PQLC2* knockout neurons compared to non-targeting control. **f,** Functional enrichment of DEGs in *PQLC2* KO neurons. **g,** Representative western blots for targets of mTORC1 in *PQLC2* KO neurons versus NTC. **h-k,**

Quantification of western blots showing that *PQLC2* knockout neurons at day 14 do not exhibit differences in phosphorylation of mTORC1 substrates compared to NTC. Throughout, n=3-5 independent experiments as indicated, * $p < 0.05$, ** $p < 0.01$, *** $p < 0.001$.
