## Extended Data Fig. 5 for "A neuronal CRISPRi screen identifies PQLC2 as a lysosomal pH regulator controlling tau homeostasis"

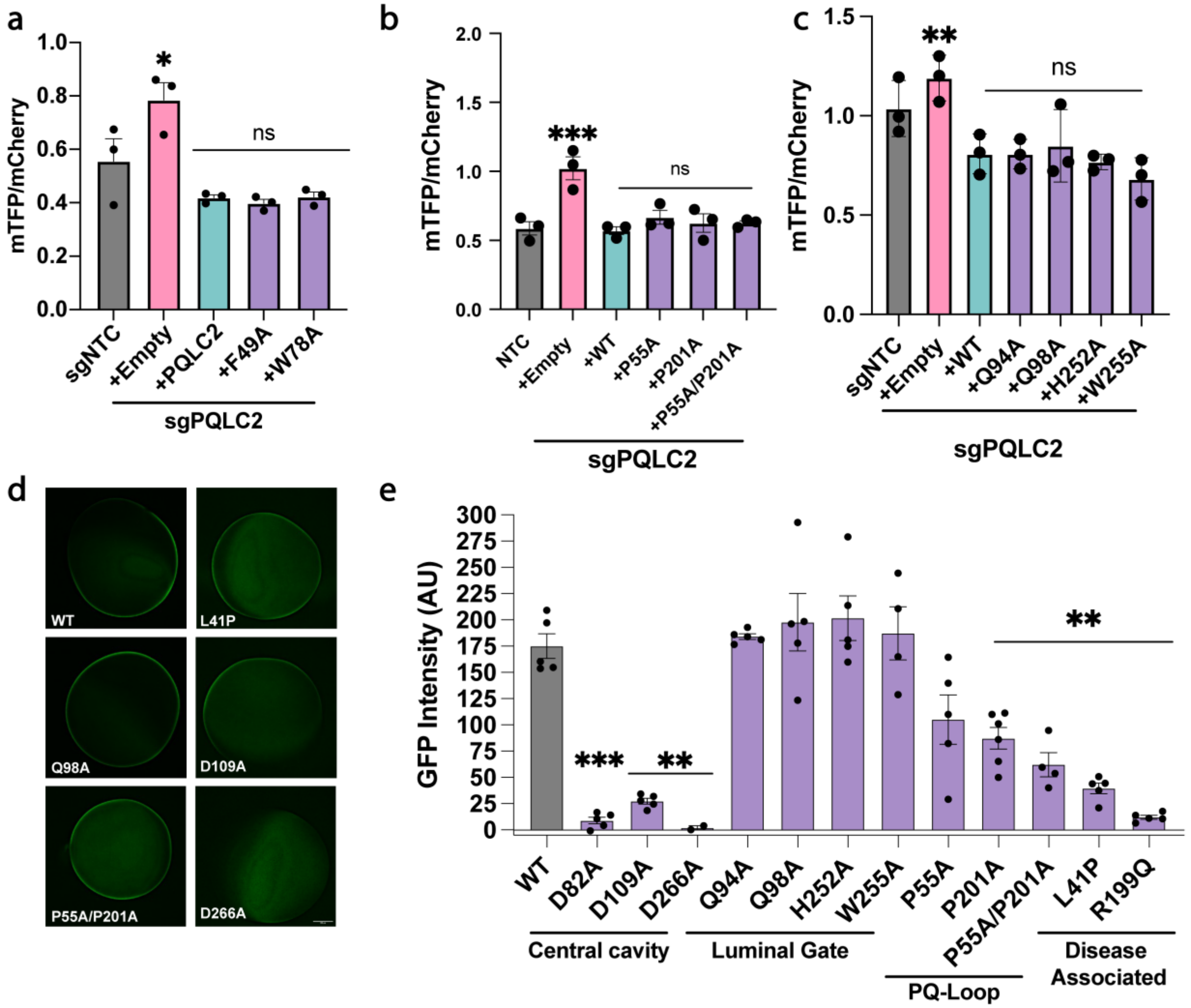


**Extended Data Figure 5: CSW complex binding, PQ-loop residues, and luminal gate residues are not required for PQLC2-mediated lysosomal pH regulation.** **a**, Expression of PQLC2 mutants that disrupt binding to the CSW complex remains sufficient to rescue lysosomal pH in Day 14 *PQLC2* CRISPRi knockdown neurons. **b**, Expression of PQ-loop mutants rescues lysosomal pH in Day 14 *PQLC2* CRISPRi knockdown neurons. **c**, Expression of amino acid mutants comprising the luminal gate of PQLC2 rescues lysosomal pH in Day 14 *PQLC2* CRISPRi knockdown neurons **d**, Representative images of *Xenopus* oocytes expressing GFP-tagged PQLC2 mutants (scale bar = 200 μm). **e**, Relative expression levels of PQLC2 mutants used in the *Xenopus* oocyte arginine uptake assay (n = 3–6 individual oocytes per mutant) asterisks denote comparison to wild-type *PQLC2* expression. Unless otherwise indicated, data represent n = 3–5 independent experiments. One-way ANOVA. * $p < 0.05$, ** $p < 0.01$, *** $p < 0.001$. Asterisks denote comparisons with the non-targeting control unless otherwise noted.
